# Parallax error indicates simple cue-anchoring in the head-direction system

**DOI:** 10.1101/2025.04.25.650191

**Authors:** Sven Krausse, Adrian J. Duszkiewicz, Emre Neftci, Friedrich T. Sommer, Alpha Renner

## Abstract

The rodent head-direction (HD) system provides an allocentric orientation signal for spatial navigation through integration of self-motion inputs and visual landmarks (‘cues’). Inferring HD from nearby visual cues faces a fundamental challenge: the direction towards the cue shifts with the animal’s position. If left uncorrected, this introduces a position-dependent parallax error. Here, we show that the HD signal in freely moving mice indeed exhibits a position-dependent bias consistent with parallax in a single-cue environment. This bias is smaller than predicted by geometric parallax and is further reduced in more natural multi-cue settings. Computational modeling revealed that this reduction of parallactic error can be explained by two averaging operations: multi-view averaging - temporal integration of the same cue viewed from different locations, and multi-cue averaging - concurrent weighting of multiple cues. Thus, averaging without explicit position-dependent correction provides a fast and robust heuristic that the HD system seems to employ for the maintenance of the internal compass. Our findings have broader implications for biological and artificial navigation systems. First, accurate allocentric reference frames are a key component of the cognitive map postulated in the extended hippocampal circuit and used for spatial cognition as well as abstract mental manipulation. Second, the employment of a fast heuristic in the HD system echoes heuristic strategies observed in other behaviors, such as decision making. More broadly, these findings highlight a trade-off in neural coding between computational efficiency and positional accuracy.

## Main

Accurate spatial navigation depends on a reliable sense of direction. The head-direction (HD) circuit is a canonical component of the brain’s spatial positioning system across diverse animal lineages (Rank, 1984; Taube et al., 1990a; Kim & Maguire, 2019; Petrucco et al., 2023). In mammals, it consists of HD cells — neurons that fire when the animal faces a specific direction in the global (i.e., allocentric) reference frame, and whose collective activity constitutes an internal compass (Taube, 2007). This compass is crucial for the maintenance of downstream spatial codes, including grid cells in the medial entorhinal cortex (Winter et al., 2015) and hippocampal place cells (Harland et al., 2017).

The HD signal originates in the dorsal tegmental nucleus (DTN) and lateral mammillary nucleus (LMN). It is relayed through the anterior thalamic nucleus (ADN) (Mizumori & Williams, 1993; Taube, 1995) to the postsubiculum (PoSub) (Rank, 1984; Taube et al., 1990b; Duszkiewicz et al., 2025) to be broadcast to higher-order cortical navigation systems. Angular head velocity (AHV) inputs update the HD signal through temporal integration (McNaughton et al., 1991, 2006; Taube, 2007; Bassett & Taube, 2001; Sharp et al., 2001). However, because AHV integration alone accumulates drift, stable orientation necessitates a mechanism for anchoring to visual cues (Taube et al., 1990a; Goodridge et al., 1998; Knierim et al., 1995; Zugaro, Berthoz, & Wiener, 2001; Zugaro et al., 2003; Raudies et al., 2015). As visual inputs are inherently egocentric, they must first be transformed into an allocentric reference frame (Skaggs et al., 1994; Zhang, 1996; Hahnloser, 2003; Bicanski & Burgess, 2016; Boucheny et al., 2005). This transformation introduces a challenge: nearby cues shift in egocentric bearing from the animal’s perspective as the animal moves, even when keeping a constant head direction, a phenomenon known as the parallax effect. This should impose a systematic, position-dependent HD bias unless corrected. To date, it remains unresolved how the HD circuit integrates egocentric visual cues to maintain a stable allocentric reference frame.

Prevailing models of the HD system either disregard the parallax effect by assuming landmarks (‘cues’) are infinitely distant (Skaggs et al., 1994; Zhang, 1996; Hahnloser, 2003; Boucheny et al., 2005), or propose explicit position-dependent corrections via hippocampal place cells (Bicanski & Burgess, 2016). The latter potentially introduces a circular dependency, as place cells are thought to rely on a stable HD signal (Mc-Naughton et al., 2006; Burak & Fiete, 2009). Thus, despite decades of research and strong evidence for cue anchoring (Taube et al., 1990a; Goodridge et al., 1998; Knierim et al., 1995; Zugaro, Berthoz, & Wiener, 2001; Zugaro et al., 2003), it remains unknown how the HD system anchors to finitedistance cues without introducing parallax errors.

Here, we show that the PoSub HD signal indeed exhibits a systematic, position-dependent bias, consistent with parallax, when anchored to a single visual cue. The magnitude of this HD bias was smaller than predicted by simple geometry. Modeling shows that parallax can be strongly attenuated via two passive mechanisms, without requiring active position-dependent correction: first, multi-view averaging, in which temporal integration of the same cue across different locations reduces position-dependent HD bias; and second, multicue averaging, in which concurrent weighting of several cues causes their individual parallax biases to cancel each other out. Importantly, the spatial pattern of the parallax-induced bias can be informative about the assumed anchoring angle between the egocentric cue bearing and head direction.

## Results

### HD signal exhibits parallax error in a single-cue environment

We first tested whether the head-direction (HD) signal is distorted by parallax in a single-cue environment. We recorded large populations of PoSub HD cells while mice freely explored an elevated circular platform (n = 823 HD cells; 11 sessions from 10 mice) and tracked the animal’s head direction using high-definition motion capture (Methods, Duszkiewicz et al. (2024)). The platform was placed inside a darkened enclosure designed to minimize the availability of uncontrolled visual cues. The only light source within the enclosure was a dim 2D LED cue mounted on one wall, visible from the platform but inaccessible to the animal (Fig. 1a, see Methods).

**Figure 1.**
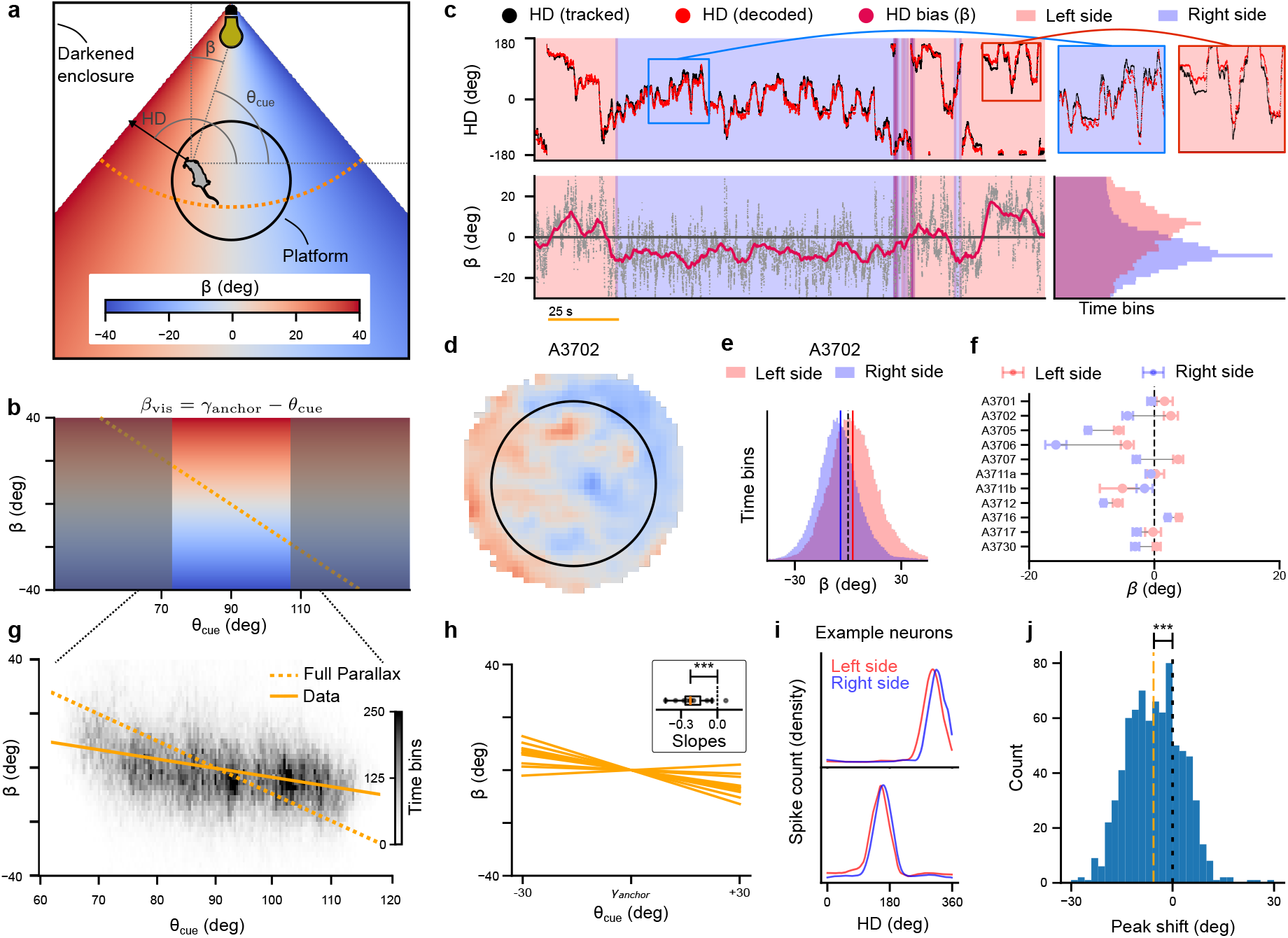
HD signal exhibits position-dependent parallax errors. (**a**) Diagram of the experimental setup. Parallax-induced HD bias β color-coded at different locations in the arena. Cue azimuth θ_cue_ and expected HD bias from parallax β are indicated in gray. Orange curve depicts linear change in the expected HD bias, as shown in panel (b). (**b**) Expected linear relationship between the parallax-induced HD bias β and the cue azimuth θ_cue_. Equation at the top can be used for linear regression to estimate the strength of the parallax effect via linear regression (panel (g)), using the γ_anchor_ = 90° as the offset between the two angles. (**c**) Example session from Duszkiewicz et al. (2024) showing the parallax effect using Bayesian decoding. Top panel shows tracked (black) and decoded (red) HD. Inset plots on the right show a zoomed-in version of two examples where the systematic HD bias (difference between tracked and decoded HD) is visible. Bottom panel shows the difference between the tracked and decoded HD. The background of the plot is color-coded to indicate whether the animal is towards the right (blue) or left (red) side of the cue. The vertical histogram on the bottom right shows the distribution of recorded HD bias when the animal is on the right/left side of the cue. (**d**) Heat map of the measured HD bias β across the platform. Color indicates the mean recorded HD bias for each spatial bin in the environment (same color map as in panel (a)). (**e**) Histogram of measured HD bias in a single recording session. The two histograms correspond to time points where the animal is either on the left (red) or right (blue) side of the circular platform. (**f**) Mean HD bias for each recording session when the animal is on the left/right side of the cue. Error bars indicate the error on the mean. (**g**) Measured HD bias vs. cue azimuth θ_cue_. Linear regression line is shown in orange and shows a significantly non-zero slope (bootstrapped linear regression, slope:− 0.35 ± 0.02, p-value < 0.001). (**h**) Linear regression lines for all recording sessions. Inset shows a box plot of the distribution of computed linear regression slopes. Overall, the slopes are significantly below zero (t-test, *t*(10) = − 4.83, *p* = 3.5 *·* 10^−4^, Cohen’s *d* = − 1.45), as expected if the parallax effect is present. (**i**) Tuning curves of two example neurons. Tuning curves are computed on two epochs: when the animal is either on the left (red) or right (blue) side of the cue. A shift in the preferred direction is detectable on a single neuron level. (**j**) Distribution of preferred direction shifts for all neurons of all sessions. The change in preferred direction was computed as the peak in the cross-correlation between the two tuning curves shown in panel (i) for each neuron in each session. The resulting distribution shows that the neurons have a significant tendency to be negatively shifted (t-test, t(822)=5.87, *p* < 1 *·* 10^−6^, Cohen’s *d* = − 0.20), indicating a parallax effect that can be measured on the single neuron level.

If the HD system fully corrects for parallax in this paradigm, the decoded HD should be independent of the animal’s location on the platform. In contrast, if the HD signal is subject to parallax, HD should be offset depending on the cue azimuth θ_cue_ (Fig. 1a,b). To disambiguate between these two possibilities, we decoded the internal HD signal from PoSub spiking activity with a Bayesian decoder (Methods) and computed the HD bias β_dec_ as the difference between the decoded HD and the real HD in the allocentric reference frame obtained from motion capture (Fig. 1c). Surprisingly, we observed a systematic dependence of HD bias on cue azimuth, consistent with parallax (Fig. 1d). When the animal was on the right side of the platform, we observed a negative HD bias, whereas positions on the left side showed positive bias (Fig. 1c-f). All but one of the animals (9/11 sessions) exhibited highly significant differences in mean HD bias between left and right conditions (Fig. 1f; Welch’s *t*-tests, all significant sessions *p* ≤ 0.003; Suppl. Tab. 1). In all significant sessions, the HD bias was positive when on the left side of the platform compared to negative on the right (Cohen’s *d* = 0.11 to 1.00, median |*d*| = 0.32), indicating that the HD signal is subject to parallax error in a single-cue environment.

To quantify the amount of parallax, we used linear regression between the HD bias β_dec_ and the cue azimuth θ_cue_ (Fig. 1c, Equations (3)-(6) in Methods, see also Suppl. Sec. Cue bearing vs. Cue azimuth). In this framework, the slope of the regression (*w*) denotes the amount of parallax. *w* = − 1 corresponds to maximal parallax, i.e., purely visual cue-based orientation, and *w* = 0 to full parallax correction. We term the x-intercept of the linear regression the *anchoring angle* (γ_anchor_) — the direction that defines the alignment of the HD signal with respect to the cue. This cue–HD alignment is also implied by cue-control experiments, in which rotating a salient cue rotates preferred directions of HD cells by an equivalent offset (Taube et al., 1990a; Goodridge et al., 1998; Zugaro et al., 2003). To account for the possibility of different anchoring angles across animals and sessions, the fit was performed separately for each session (Fig. 1h). We observed fitted slopes significantly below zero in 10 out of 11 recording sessions (mean slope 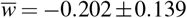), meaning that the parallax effect was apparent in all but one session (boot-strapped t-test, p-values < 0.05, Methods, Suppl. Tab. 4). However, the maximal negative measured slope was − 0.43, indicating a lower position dependence of the systematic error than the *w* = − 1 predicted by purely visual cue-based orientation. To exclude the possibility that these findings on the population level are an artifact resulting from biases in the tuning curves of a few neurons, we repeated the analysis while constraining the tuning curve estimation to time intervals where the animal was in the center of the environment. All findings presented above could qualitatively be reproduced (see Suppl. Sec. Analysis of tuning curves from only the center of the environment).

Given that the parallax error was reliably detected at the population level, we next asked whether it is also expressed at the level of individual HD cells. Establishing this is important because a population-level bias could, in principle, be driven by a small subgroup of neurons with idiosyncratic tuning, or by decoding-related artifacts. If, however, parallax affects the tuning of single HD neurons, their preferred firing direction should shift systematically with the cue azimuth experienced by the animal. To test this, we split the sessions into two conditions depending on whether the animal was on the left or right side of the platform. We computed HD tuning curves separately for left and right conditions (Fig. 1i) and quantified the shift between the two tuning curves using cross-correlation. We observed a significant clockwise shift of the left-side tuning curves relative to the right-side (mean = −4.78° ± 0.81°, *n* = 823 neurons, t(822)=5.87, *p* < 1 × 10^−6^, Cohen’s *d* = − 0.20). The distribution of peak shifts across the full population shown in Fig. 1j indicates that parallax is not restricted to a few outlier cells but instead represents a robust, system-wide property of the HD network.

### Parallax errors reveal the anchoring angle

Cue-control experiments show that salient cues can quickly gain control over the head-direction (HD) signal, implying that the system learns a fixed alignment between egocentric cue bearing and its internal compass (Taube et al., 1990b; Goodridge et al., 1998; Zugaro et al., 2003). The *anchoring angle*, γ_anchor_, introduced above, captures this alignment. Defined as the x-intercept of the bias-azimuth regression, it marks the cue azimuth at which the parallax errors vanish. This makes the spatial pattern of parallax errors a useful measurement tool for estimating how the HD representation is anchored to visual cues. Intuitively, γ_anchor_ can be thought of as the head direction at which cue control was first established, i.e., the *anchoring event*. When the cue reappears after a period without it, the system should tend to correct its internal compass back towards this offset.

To estimate γ_anchor_ from the data, we examined the spatial distribution of parallax errors across animals. We observed distinct spatial patterns of HD bias in individual recordings (Fig. 2a, Suppl. Fig. 13), suggesting that γ_anchor_ is session-specific. We quantified this by fitting the linear regression between HD bias and cue azimuth and identifying the cue azimuth at which the HD bias crossed zero (the x-intercept). Because this estimate is very sensitive to noise, we restricted the analysis to sessions with consistent linear relationships (standard deviation of slope *w* < 30°, *n* = 7 sessions). The resulting γ_anchor_ estimates were stable within sessions, showing agreement between the first and second halves of each recording (Fig. 2b). Across sessions, γ_anchor_ clustered within about 40° of the platform center, consistent with anchoring occurring when the cue lies within the mouse’s field of high visual acuity (van Beest et al., 2021).

**Figure 2.**
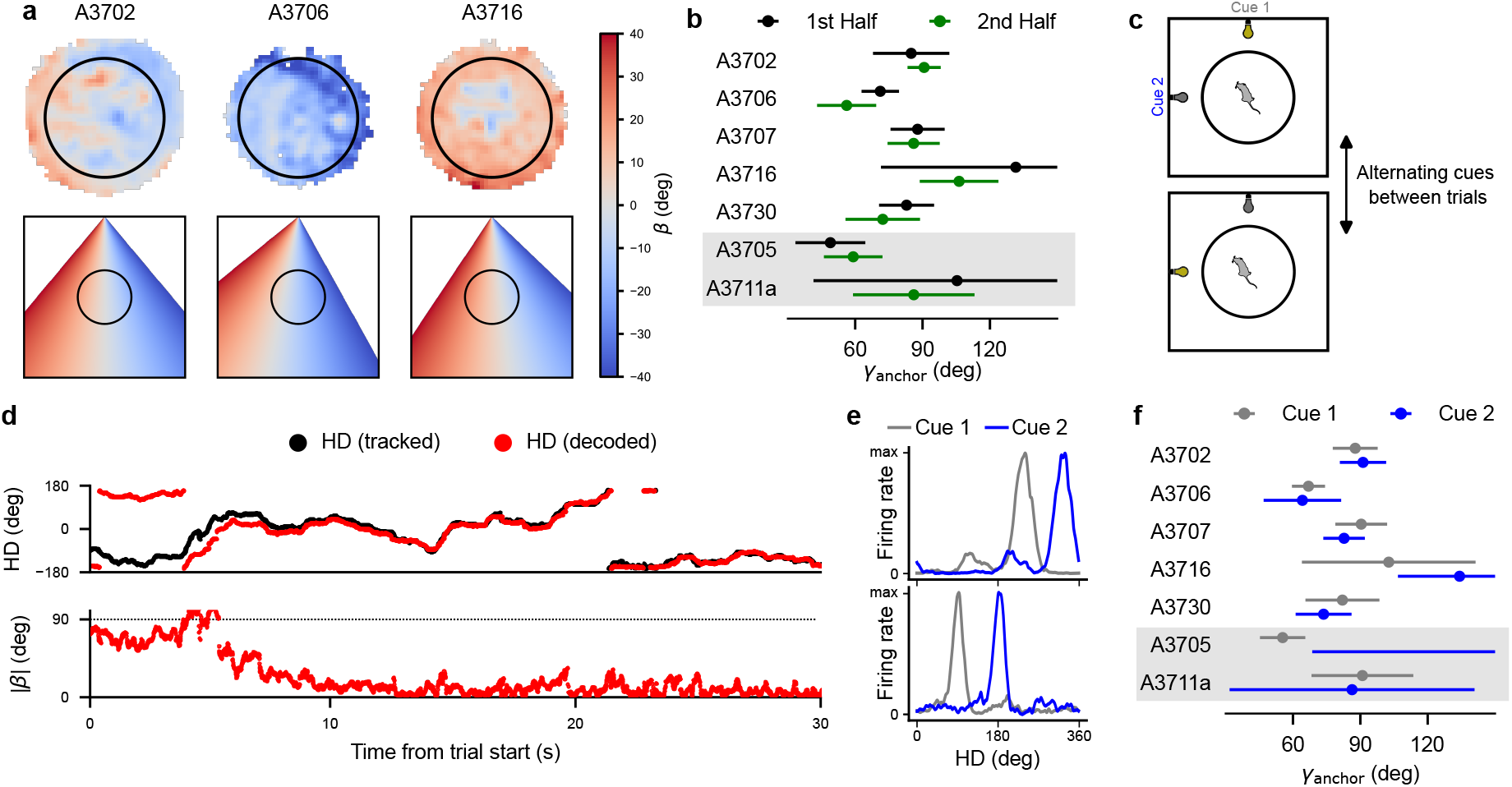
Estimating the anchoring angle γ_anchor_ based on parallax effects. (**a**): Top panels: Heat maps of the animal during three different recording sessions, color-coded by the observed angular error β_dec_ of the neural data at each location. Bottom panels: Theoretical predictions of the spatial distribution of β from a purely vision-based HD model. (**b**): Anchoring angles for included sessions from the single-cue dataset, as computed from the linear regression results (Eq. 7). Anchoring angles are computed for the first and second halves (split over time) separately. Note that the first five sessions in this plot were sessions that exhibited reorientation after the cue switch occurred, while the bottom two sessions (*A3705* and *A3711A*, grey background) did not show reorientation. (**c**): Illustration of cue switching between different trials of a recording session. Between each trial, the position of the single cue was changed to the center of a different wall, then throughout the recordings, the two cues were presented alternatingly over the course of 16 trials (8 per cue). (**d**): Example trace of the first 30 seconds after a trial start. After an adaptation period, the decoded HD shifts to the 90-degree rotated cue. (**e**): Two example tuning curves of a neuron in the two different cue conditions. The preferred firing direction shifts in the two different cue conditions by the amount of rotation of the cue (90 degrees). (**f**): Anchoring angles computed for the seven sessions in the two different cue conditions. The anchoring angles for the original and rotated cues indicate that the anchoring angle remains relatively stable over time and across rotations of the cue.

We further tested the stability of γ_anchor_ during cue rotations (Fig. 2c). In each recording session, after the initial period with a stable cue, the location of the LED cue was repeatedly switched between two adjacent walls of the enclosure. In sessions where the HD system reliably realigned to the rotated cue (Fig. 2d,e), the estimated γ_anchor_ before and after rotation overlapped within confidence intervals (Fig. 2f, top, *n* = 5). This is consistent with the view that the system treated the rotated cue as the same landmark, maintaining the same anchoring angle. In contrast, in sessions where realignment to the other cue location failed (*n* = 2), γ_anchor_ could not be estimated reliably (Fig. 2f, grey), consistent with the system treating the rotated cue as novel and unanchored. Together, these findings show that the anchoring angle can be inferred from the spatial structure of parallax errors, indicating that HD anchoring occurs at a specific bearing relative to visual cues.

### Parallax errors are attenuated in a standard recording environment

In natural settings and most laboratory experiments, animals are surrounded by multiple cues, even if a single cue may dominate the visual field of view. In such conditions, the parallax errors introduced by individual cues could average out, reducing the overall distortion without the need for active correction, which might explain why parallax errors were not observed in previous studies of the HD system (Taube et al., 1990b).

We therefore analyzed additional PoSub recordings from animals foraging in a standard square open-field environment with a single cue card (n = 1041 HD cells; 16 sessions from 16 mice) (Fig. 3a). In contrast to the single-cue paradigm, this environment was illuminated, thereby providing several additional unintentional cues, likely including the corners and boundaries of the enclosure. We found that while we still observed a parallax effect relative to the cue card when the cue card was in view, its magnitude was substantially reduced compared to the single-cue paradigm (Fig. 3b-d; Welch’s t-test, p-value < 0.001, Suppl. Tab. 3).

**Figure 3.**
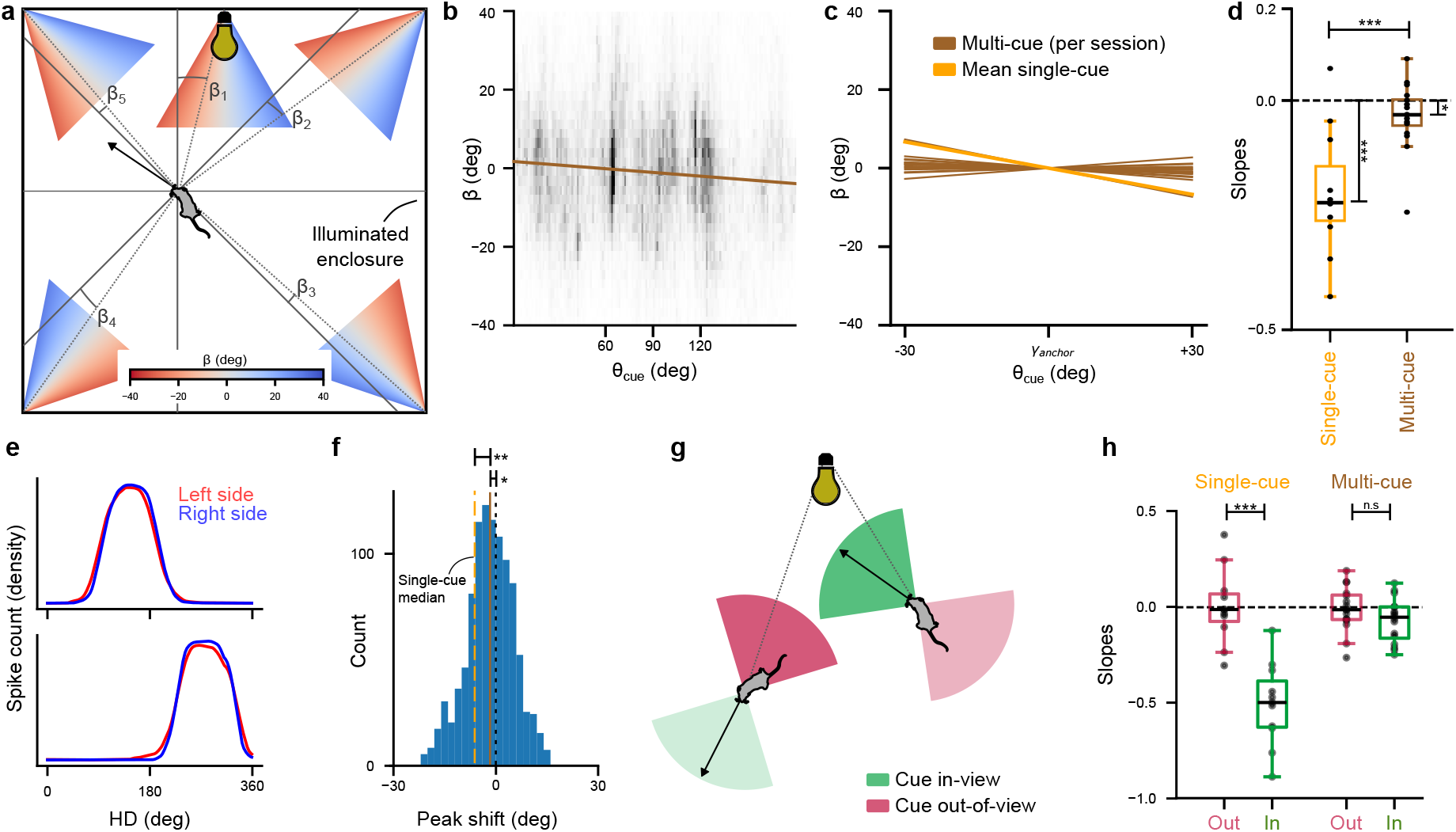
Reduced parallax error in a multi-cue environment. **(a)**: Diagram of the experimental setup in the multi-cue environment. The animal is navigating in an illuminated environment with a fixed cue on the top wall. Additional features in the room can additionally be used as anchors for the HD system and should therefore have their own parallax error, illustrated by the different color maps. The choice of the four corners as additional cues is merely illustrative - in principle, any additional visual cues could be used to anchor the HD. **(b)**: Linear regression performed on a representative session. Relationship between the azimuth to the cue card θ_cue_ and the observed HD bias β_dec_. Individual data points are calculated from the decoded HD traces; the brown line indicates the fitted linear regression. The range of θ_cue_ is determined by the size of the environment. **(c)**: Linear regression lines for all 16 recording sessions in the multi-cue environment (brown lines). For comparison, we also show the median regression line from the single cue environment (Fig. 1h). **(d)**: Box plot for measured slopes in the single cue and multi-cue environments. The means of the slopes are significantly below zero in both the single cue and multi-cue setting (t-test, single cue *p* < 10^−3^, multi-cue *p* < 0.05, Suppl. table 2). Slopes in the single cue dataset are significantly more negative than in the multi-cue dataset (t-test, *p* < 10^−3^, Suppl. table 3). **(e)**: Example tuning curves in multi-cue dataset, split by the animal position in the arena (left - red, right - blue, compare Fig. 1i). **(f)**: Distribution of shifts in tuning curves when the animal is on the left/right side of the platform. Data aggregated over all HD neurons in all 16 sessions. For comparison, we also show the median peak shift of the single cue dataset analysis (orange dotted line, Fig. 1j), which is significantly more shifted than the multi-cue median peak shift (t-test, *p* = 2.1 · 10^−4^). **(g)**: Sketch to illustrate the ‘in-view’ (green) and ‘out-of-view’ (red) conditions. Vision cones filter the data for time periods where the cue is right in front (in-view) or behind (out-of-view) the animal. **(h)**: Resulting slope as an indicator of parallax strength in the in-view (green) and out-of-view (red) conditions for the single cue and multi-cue data. Significant differences are observed in the single cue dataset (t-test, *p* < 10^−3^), whereas the slope remains close to zero in the multi-cue dataset in both conditions (t-test, *p* = 0.165, Suppl. table 3).

We also performed the single neuron tuning curve analysis for this multi-cue data (Fig. 3e,f). The shift between tuning curves was weaker than in the single-cue paradigm but still significant (mean = − 1.17° ± 0.58°, *n* = 1041 neurons,, *t*-test, *t*(1040) = − 2.02, *p* = 0.044). The shift magnitude differs significantly between the experimental paradigms (two-sample *t*-test, *t*(1862) = 3.71, *p* = 2.1 × 10^−4^), confirming the results of the system-level analysis, that multiple cues help correct for the parallax error from each individual cue.

### The parallax effect is dependent on cue visibility

In addition to using visual cues, animals rely on integrating self-motion information, which is not subject to parallax. This is most apparent during time intervals with no available visual cues, in which the animal must rely solely on internal self-motion information like angular head velocity (AHV). We therefore investigated whether the observed parallax effect disappears when the visual cue is not within the mouse’s field of view (out-of-view condition) in comparison with periods where the animal is directly facing the cue (in-view condition) (Fig. 3g). We expected to see a difference between the two conditions in the single-cue but not the multi-cue paradigm, as the animal could use additional cues for visual position estimation in the multi-cue paradigm, even when the intentionally placed cue card was out of view.

We indeed observed lower HD bias when the cue was behind the mouse and could therefore be assumed to be out of view. In the single-cue paradigm, 10 out of 11 sessions showed a significantly lower regression slope in the out-of-view condition (Suppl. Tab. 4) than when the animal was facing the cue directly (in-view) (t-test p-value < 0.001, Fig. 3h, Suppl. Tab. 3). As expected, the multi-cue paradigm did not show a significant shift between the in-view and out-of-view conditions (Fig. 3h, Suppl. Tab. 3), indicating that parallax depends on the visibility of a single dominant cue and is effectively suppressed when multiple cues contribute to the HD estimate.

### Simple integration passively attenuates parallax error

The observed attenuation of the parallax error could, in principle, arise from an active position-dependent mechanism (Bicanski & Burgess, 2016). However, it may also emerge passively through two simpler processes: *Multi-view integration*, where the animal sees the cue from many different viewpoints and integrates these over time, and *multi-cue integration*, where cues with different parallax average out. To assess whether the simplest mechanism, basic cue anchoring, as suggested by cue-control experiments and our data, is sufficient to attenuate parallax error, we next provide a minimal model of internal HD generation without any position-dependent corrections.

The HD system is thought to integrate AHV from vestibular and proprioceptive inputs while using visual cues to prevent drift (Blair & Sharp, 1996; McNaughton et al., 1991; Goodridge et al., 1998; Knierim et al., 1995; Zugaro, Berthoz, & Wiener, 2001; Zugaro, Tabuchi, et al., 2001; Bicanski & Burgess, 2016; Ocko et al., 2018; Page & Jeffery, 2018). Integrating these two signals naturally attenuates parallax, because anchoring to the visual cue occurs at varying cue azimuths and therefore averages over time. This averaging effect is expected to be even stronger with multiple cues (Gaussier et al., 2019), making complex correction mechanisms potentially unnecessary.

To test this quantitatively, we implemented a simple integrate-and-vision model (Fig. 4a), in which the HD estimate α_iv_ is a weighted sum of the AHV-integrated estimate α_int_ and the vision-based estimate α_vis_:

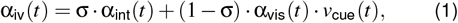

where σ is the relative weighting of AHV integration and visual anchoring and *v*_cue_(*t*) is a binary variable indicating whether the visual cue is within the animal’s field of vision. The integrated estimate evolves as

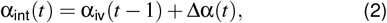

with the angular head displacement ∆α(*t*) derived from the head tracking data. The “simulated mice” follow the same path and orientation as the real animals from the neural recordings.

**Figure 4.**
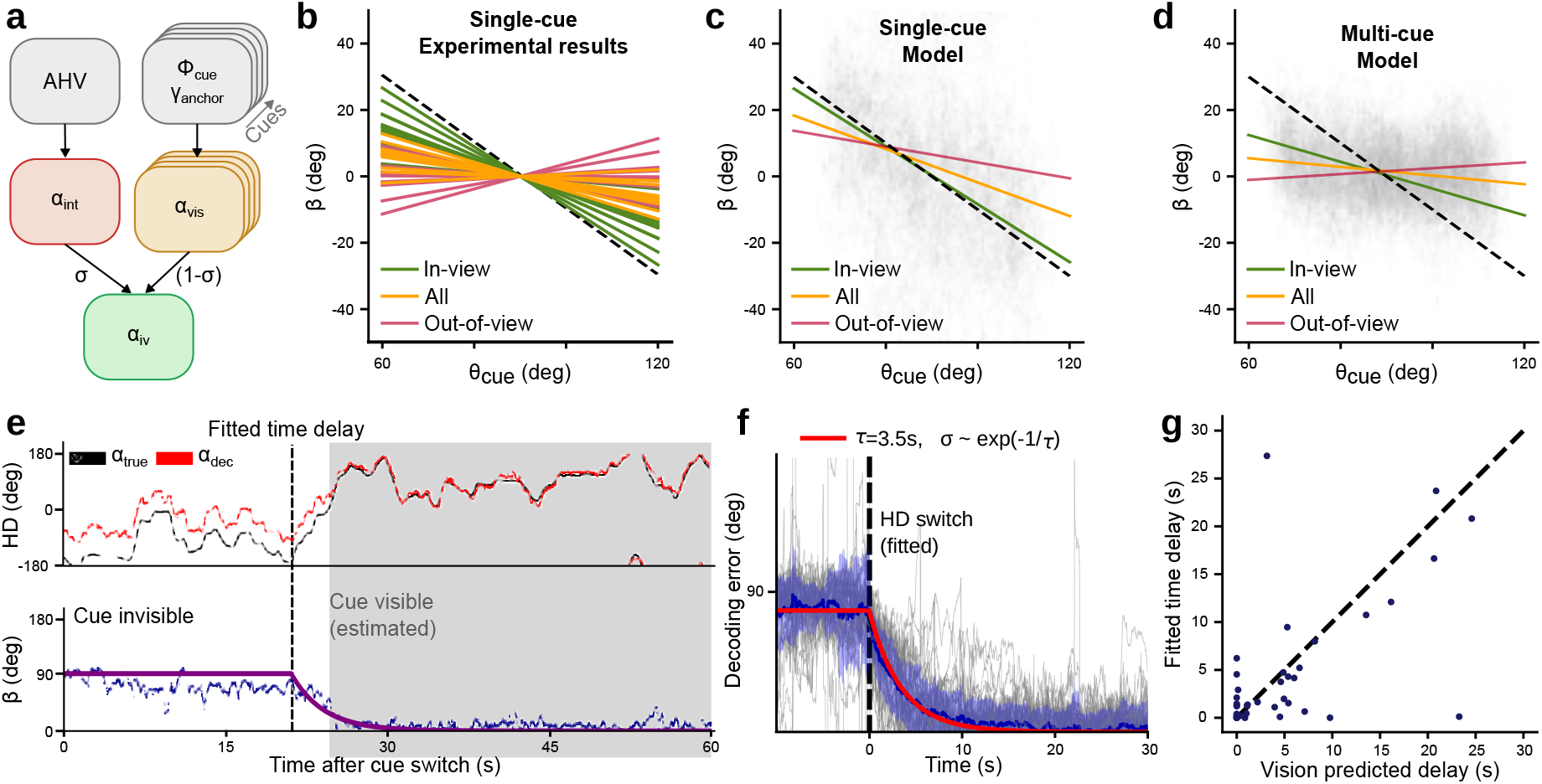
Simulations confirm that AHV and the use of multiple cues neutralize the measured parallax effect. (**a**): Visualization of the simulated HD model, as described in Eqs. (1, 2). (**b**): Single-cue experiment experimentally determined linear regression slopes as a reference for the modeling results in panels (c) and (d). Slopes for complete sessions are shown in orange, filtered for the in-view condition in green, and out-of-view condition in red. (**c**): Fitted slopes from simulated head direction using a single cue. A negative slope with a magnitude below 1 (black dashed line) is observed in all three conditions. The slope is steeper in periods when the cue is in view (green) and less steep when the cue is out of view (brown), matching our experimental observations. (**d**): Parallax analysis on the simulated data with multiple cues (cue card and the four corners of the square environment). Overall, parallax is drastically attenuated compared to (b). (**e**): Example beginning of a trial with fitted HD switching onset and exponential decay fit shown in the bottom panel. Gray background color indicates when the cue is in-view, according to a 180° field of vision. (**f**): All HD reorientation events aligned on the fitted time delay. The resulting mean decay shows a time constant τ = 3.5*s*, which we use to determine the weighting between visual and integrated HD estimates. (**g**): Distribution of fitted time delays and time delays predicted by a 180° field of vision.

The model has two free parameters: σ and the angle of the field of vision. Both were estimated from reorientation events following cue rotation in the single-cue dataset. For each reorientation event, we computed the decoding error relative to its final value post-reorientation. During reorientation events, we observed exponential decay of the decoding error from the initial value of 90° towards 0° (Fig. 4e). We fit the onset of this reorientation event and the decay constant of the exponential decay across all reorientation events (see Methods). As the decay constant τ indicates how strong the influence of the new cue position is, it is directly related to the weighting parameter σ in our model (Fig. 4f), resulting in an estimated weighting parameter σ = 0.997. We additionally used the fitted delay until the onset of the reorientation event to define the effective vision cone of the animal. Intuitively, the cue has to enter the animal’s field of vision before the reorientation is initiated. We tuned the effective field of vision so that the fitted onset best aligns with the time points where the cue in the new position appears in the animal’s view, resulting in an effective cone of vision of 180° (Fig. 4g).

The simplified integrate-and-vision model is intended to demonstrate the principle of passive parallax attenuation, not to capture the full complexity of multi-sensory integration. In reality, the weighting between AHV integration and visual input likely varies with cue familiarity (Cheng et al., 2007; Julian et al., 2018) and the uncertainty (or variance) of α_int_ and α_vis_, which could explain phenomena such as distance-dependent remapping (Zugaro, Berthoz, & Wiener, 2001). Because the variance of α_int_ in the neural system is unknown and AHV derived from the tracking system is very precise, we used a fixed σ and added a small amount of Gaussian noise to the AHV input (mean 0 °/s, s.d. 5 °/s) to obtain more realistic variability.

Repeating the linear regression parallax analysis on the modeled HD (treating α_iv_ as internal HD) confirmed that AHV integration is sufficient to reduce parallax relative to the geometric prediction of − 1 for a purely vision-based system (Fig. 4b, black dashed vs. orange). The modeled slope *w* was sensitive to σ, which we cannot estimate with high precision, but the qualitative effect was robust. The model also reproduced the difference between cue-visible and cue-invisible conditions (Fig. 4c, green vs. red): slopes were steeper when the cue was visible. As expected, the inclusion of AHV integration also improved the HD tracking accuracy compared to a purely vision-based model, which would be subject to the full parallax effect (Suppl. Fig. 8).

We next extended the model to multiple cues. For each visible cue, we computed α_vis_ (Eq. 3) and took the circular mean across cues, using equal weighting for simplicity. Additional cues were assumed at the four corners of the enclosure. In this multi-cue setting, the parallax bias almost vanished (Fig. 4d).

This simple integrate-and-vision model therefore demonstrates that the position-dependent parallax bias expected from geometry does arise in a single-cue environment, but is passively attenuated by AHV integration, and is almost eliminated when multiple cues are present. No active correction mechanism is required, consistent with the experimental findings above.

## Discussion

Spatial navigation requires a stable allocentric sense of direction, yet sensory cues are perceived egocentrically. This necessitates a coordinate transformation whose underlying neural mechanism has been a long-standing question (Taube, 2007; Zhang, 1996). In this study, we provide evidence for a systematic parallax error in the HD signal of mice navigating an environment with only a single visual cue. This finding indicates that the HD system does not employ a complex, position-dependent correction mechanism, for instance, through place cell feedback, which would introduce a circular dependency, as place cells themselves require a stable HD signal (Bicanski & Burgess, 2016; McNaughton et al., 2006). Instead, our results support a model where the system relies on a simple, computationally efficient egocentric cue-anchoring strategy for the ego-to-allocentric transformation (Skaggs et al., 1994; Page & Jeffery, 2018). This process, while effective, inherently introduces a predictable, position-dependent bias when referencing a single nearby cue.

The single-cue environment, although artificial, provides a unique window into the mechanism of extracting HD from sensory input. Understanding this mechanism is critical, as a stable allocentric HD signal is a fundamental prerequisite for downstream circuits in the hippocampal formation to build and maintain a coherent cognitive map (McNaughton et al., 2006; Raudies et al., 2015; Tan et al., 2017).

### A mechanism for anchoring the HD signal

The findings presented here suggest a specific model of how visual input is integrated in the HD network. A major challenge of this integration is the fact that visual observations are inherently egocentric, but the head direction is an allocentric signal. Therefore, the HD network needs to perform an egocentric-to-allocentric coordinate transform. This coordinate transform is widely believed to involve the retrosplenial cortex (RSC) and PoSub (Goodridge & Taube, 1997; Clark et al., 2010). RSC contains cells encoding the egocentric angle with respect to specific environmental features and HD cells driven primarily by visual cues (Alexander et al., 2020; Jacob et al., 2017). PoSub thus receives input both from the visual HD pathway through the RSC, as well as from the movement-integrating pathway through the ADN (Fig. 5). For a detailed description of the movement integration pathway and how to model it, see Bicanski & Burgess (2016).

**Figure 5.**
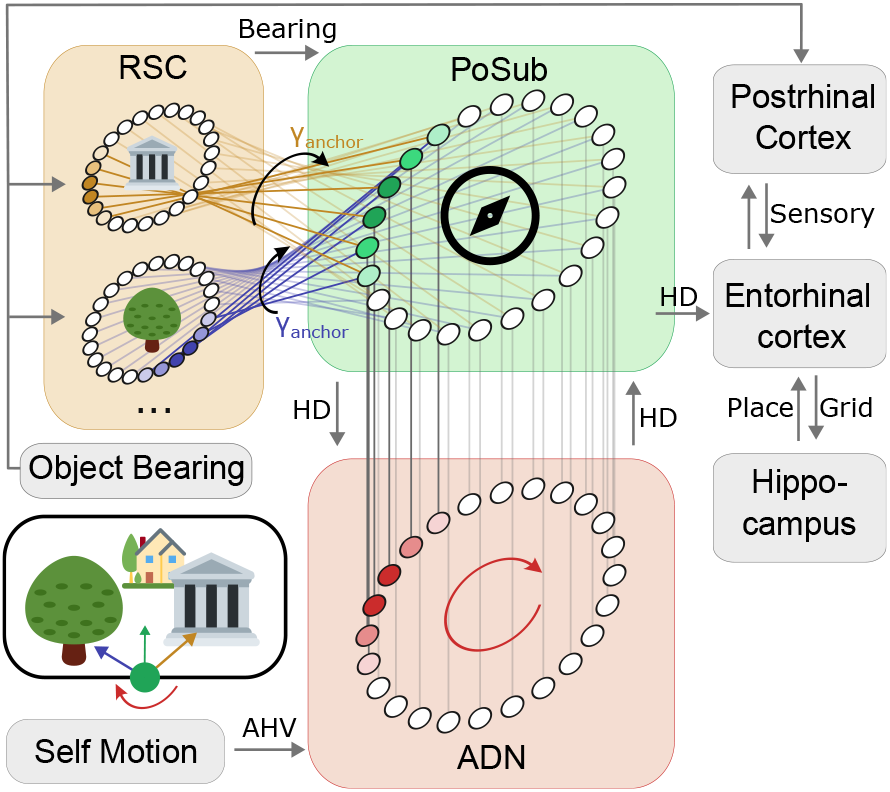
Conceptual model of the HD formation from combined AHV integration and visual inputs. Based on visual inputs and inertial movement measurements (bottom left), visual cortices and subcortical areas transmit information along different pathways. The visual information is mapped to bearing cell rings in the retrosplenial cortex (RSC). These, in turn, have a fixed anchoring angle to map onto the head direction cells in the postsubiculum (PoSub). The inertial measurements in turn are projected through LMN to the anterodorsal nucleus (ADN) (see Bicanski & Burgess (2016) for a detailed description of this mechanism). PoSub acts as a central integrator in this model, combining the visual and integrated HD states and relaying them back to RSC and AdN and projecting them forward to the entorhinal cortex for subsequent position-estimation in the hippocampal formation.

According to the current literature, the exact mechanism that allows the egocentric-to-allocentric transformation in the HD network is still unknown (Yoder et al., 2011; Jeffery et al., 2016; Duszkiewicz et al., 2025). The experimental results provided in this study suggest the following conceptual model: Bearing cells in the RSC map onto an HD ring attractor in Po-Sub with a fixed offset. This fixed offset is what determines the anchoring angle in the previous analysis. Specific bearing cell ring attractors may be activated by visual cues, whose identity and egocentric position are provided by the visual system (Fig. 5).

In this model, any single bearing cell ring attractor is subject to the parallax effect. Therefore, in a highly restricted setting of only one singular cue in the environment, this parallax error is also present in the PoSub. However, under natural conditions, multiple cues activate multiple bearing cell ring attractors. The postsubiculum then averages the HD estimates from the different visual cues together with the movement integrated input from the ADN (akin to the simulated α_iv_ above).

The weighting between the different bearing cell rings and the ADN input could be established by the gain of each population (effectively the height of the activity bump in each ring attractor) (Ajabi et al., 2023). The gain of each ring attractor may be determined by how fast the activity bump is moving along the attractor manifold, where immobile activity bumps are weighted higher than fast-moving ones. Such a weighting mechanism would result in close-by visual cues to influence the PoSub HD less, as their azimuth changes faster than those of more distant cues. This weighting would also give more weight to stationary objects rather than moving ones. It would finally also weigh movement-integrated estimates vs. visual estimates, relying more strongly on vision when the head is moved quickly, and the movement integration is more prone to inaccurate HD estimates. Finally, in our conceptual model, as already suggested in previous studies, PoSub sends its combined HD signal back to RSC and ADN/LMN as a correction mechanism. Additionally, PoSub propagates the combined HD estimate forward to the hippocampal formation for subsequent position estimation. This direct mapping strategy of selfmotion information and bearing information of multiple objects onto one ring attractor is fast and computationally efficient, as it does not require explicit positional information to compute an HD estimate. Such efficiency is a critical advantage for both biological and artificial navigation systems, where energy and processing resources are constrained (Arleo & Gerstner, 2000; Kreiser et al., 2020; Renner et al., 2022).

### The anchoring angle as a link between egocentric and allocentric frames

The anchoring angle provides a simple geometric description of how egocentric visual cues become aligned with the allocentric HD representation. It effectively marks the phase offset between the retrosplenial populations encoding the egocentric cue bearing and the postsubicular HD ring that represents allocentric head direction (Alexander et al., 2020; Bican-ski & Burgess, 2020; Lozano et al., 2017). Once established, this offset acts as a stable “reference phase” between sensory and internal frames of reference. Here, we have shown that the parallax effect can be used as a measurement tool to estimate this important property of the egocentric to allocentric mapping.

Conceptually, the anchoring angle can be viewed as a memory of the cue azimuth (the angle towards the cue in the allocentric reference frame) at which cue control was first acquired. When the same cue reappears, the HD system may tend to pull the internal compass back toward this learned offset. This explains why, after experimentally introduced cue rotations, the system realigns instantly when the cue is recognized as familiar but fails to do so when it is perceived as novel.

### Why parallax error in the HD system was not observed in previous studies

The absence of previous reports on the parallax effect warrants explanation. Detailed analysis of our data suggests two reasons. First, the HD system continuously integrates AHV, which inherently smooths the visual error over time as the animal moves through the environment (McNaughton et al., 1991; Taube, 2007). Our data directly support this, showing that the measured HD bias was significantly attenuated when the cue was not in the animal’s direct field of view. Second, and more importantly, naturalistic environments are rich with visual and other directional sensory cues. Our analysis of recordings from a multi-cue setting revealed a practical elimination of the parallax error. This strongly supports the hypothesis that the HD system averages heading estimates derived from multiple cues to form a unified, robust signal (Gaussier et al., 2019; Goodridge et al., 1998). The individual parallax errors associated with each cue, having different spatial dependencies, effectively cancel each other out. Most experimental paradigms, even those with a primary cue card, likely provide sufficient secondary cues (e.g., arena corners, arena geometry, shadows, equipment, odors, sounds) to enable this averaging, thus masking the parallax from any single cue.

### Parallax error is a system-wide phenomenon

The impact of parallax is not limited to a subset of specialized cells but is a system-wide phenomenon. Our single-neuron analysis revealed that the preferred firing directions of individual HD neurons shift systematically with the animal’s position relative to the cue. This confirms that the entire HD attractor network is biased by parallax in the single-cue context, rather than the effect being an artifact of a few bearing-like cells.

## Conclusions and future directions

In conclusion, our study demonstrates that the mammalian HD system is subject to parallax error when constrained to a single visual cue. This finding strongly supports a model of simple egocentric cue-anchoring, a computationally efficient but imperfect strategy for ego-to-allocentric transformation. The apparent vulnerability to error is robustly overcome in natural settings through the parallel integration of self-motion cues and the averaging of information from multiple environmental cues. This work clarifies a key mechanism of spatial cognition and highlights an elegant solution for generating a stable allocentric signal, offering insights that can inform the development of more robust and efficient bio-inspired navigation systems (Arleo & Gerstner, 2000; Kreiser et al., 2020; Renner et al., 2022).

Our work has broader implications. The results suggest that fast, approximate heuristics that brains use in higher-level cognition (Gigerenzer & Goldstein, 1996; Simon, 1997), often referred to as “bounded rationality”, also extend to lower-level sensory processing, such as the extraction of head direction from visual cues.

Another intriguing possibility is that our results might help investigate how brains navigate abstract mental spaces in higher-level cognition, as postulated by the cognitive map theory for the larger hippocampal network. A prerequisite for mental navigation in a cognitive map is the establishment of “allocentric” reference frames in a mental space (Behrens et al., 2018). It is open whether mental navigation uses a computation of the sort we describe, and whether even the same circuits in the postsubiculum could be involved.

Future research should build on these findings to investigate whether parallax error propagates to other nodes of the navigation circuit, such as the medial entorhinal cortex, which could further elucidate how the brain ensures a stable directional reference for navigation.

## Methods

### Single cue experiment

We analyzed neural recordings from the postsubiculum (Po-Sub) of six sessions of 6 mice during the “Cue rotation task” previously reported in (Duszkiewicz et al., 2024). We additionally included 5 previously unpublished sessions of the same experimental setup, resulting in an overall number of 11 sessions from 10 mice. Each session includes the tracked head position and direction (tracked using the Optitrack motion capture system), as well as the recorded spike trains. The mice freely explored a 30 cm diameter elevated circular platform within a 90 cm darkened square enclosure (Fig. 1a) for approximately 75 minutes per session. The recording enclosure was placed inside a metal frame, which was sealed with a black opaque cloth from all sides. Before each recording session, the recording arena was thoroughly cleaned with antibacterial wipes. Additionally, during the experiment, a strong masking odor (an air-freshener) was sprayed evenly underneath the platform. Visual cues, consisting of dim LED strips forming a “V” or “X” shape (approximately 10 cm wide), were positioned at the center of each wall and treated as point sources for analysis. During recording, all lights in the room except for the infra-red camera LEDs were switched off. Surgical procedures, electrophysiological recording details, spike sorting and neuron classification procedures, as well as the exact experimental protocol, are described in (Duszkiewicz et al., 2024). Each recording session consisted of a 1h sleep epoch followed by the cue rotation experiment. During the sleep session, the mouse was placed in its home cage on top of the circular platform. The cue rotation experiment started with a stable cue presentation (habituation period, 14-21 minutes). Subsequently, this initial cue was turned off, and a cue on an adjacent wall was activated. The experimental protocol comprised sixteen 200-second trials, with the active cue alternating between the two LEDs after each trial. Only one cue was switched on at any time. In the additionally provided 5 recording sessions, the mice’s HD neurons did not reorient during the cue rotation, but kept their orientation throughout the sessions. Unless otherwise specified, we reoriented the position data in each trial such that the cue was effectively always positioned at the north wall (Fig. 1a).

### Multi-cue experiment

We also analyzed publicly available square open field recordings of 16 mice originally described in (Duszkiewicz et al., 2024). Neural activity was again recorded from PoSub, and the animal’s position and head direction were tracked. The environment consisted of a 90 cm x 90 cm square box with a single cue card on one of the walls to serve as a cue. A key difference between this experimental setup and the previous “single-cue” environment is the lighting condition. The environment was illuminated, allowing the animal to also see the enclosure and any features within the environment clearly. Therefore, while only a single intentional cue has been added to the environment, the animals may use additional features as cues to anchor their HD estimates. Surgical procedures, electrophysiological recording details, and the exact experimental protocol are again described in (Duszkiewicz et al., 2024).

## Data Exclusion

In all datasets, we excluded periods where the mice exhibited prolonged immobility, as these periods led to less activity and an unreliable decoded HD signal. We achieved this by estimating the animal’s speed from the tracked position. We then smoothed this position signal (square kernel, 0.5 s width) and rejected time periods in which the absolute value of the smoothed speed was below 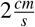 . We additionally excluded the time periods that were used to compute the tuning curves for each neuron from all further analysis. For the tuning curve estimation, we used the first 10 minutes of each recording. In the single cue experiment sessions, we also estimated the time period of HD reorientation at the beginning of the trial and excluded these time windows from the parallax analysis. In the sessions without reorientation, we only considered the trials of the original cue orientation unless otherwise specified.

## HD Decoding

As in Duszkiewicz et al. (2024), we performed Bayesian decoding (Zhang et al., 1998) on the head direction cells (cell labeling adopted from the dataset). Tuning curves for all head direction neurons were computed using the first 10 minutes of the recording (part of the habituation period in the single cue experiment; first 10 min of recording in the square arena dataset). This period was excluded from subsequent analysis. The Bayesian decoder treats these tuning curves as conditional probability distributions of head direction given a neuron’s firing. For each time bin of 10 *ms*, the likelihood distribution of the head direction was calculated by multiplying the tuning curves of all spiking neurons within that bin. We use the smoothed posterior distribution of the preceding time bin as a prior for the current time bin to discourage large jumps between consecutive estimates. The head direction estimate for each time bin was then obtained by identifying the peak of the resulting posterior. Notably, the number of HD neurons per dataset varied between all sessions (25 − 117 recorded HD units, mean: 69 ± 23 units). This led to a difference in decoding quality (mean RMSE: 19.8° ± 7.3°). RMSE and number of neurons for all sessions can be found in Suppl. Fig. 14.

### Cue visibility conditions

In Sec. The parallax effect is dependent on cue visibility, we test whether the visibility of the cue influences the amount of parallax detected in the experimental data. We differentiate between an “in-view” and “out-of-view” condition and perform the parallax analysis on both separately. For the “in-view” condition, we only consider time periods in which the egocentric viewing angle ϕ_cue_(*t*) is within ± 45°, meaning that the cue is within a 90° vision cone of the animal. The “out-of-view condition” is likewise defined as time periods in which ϕ_cue_(*t*) is within 180° ± 45°. We use these rather small cones of vision as a conservative measure to ensure that the animal is facing either directly towards or away from the cue. The data rejection criteria are also applied to these two subsets of the data before any further analysis.

### Linear regression and bootstrapping

In a simple cue anchoring model, there exists a fixed mapping between the egocentric cue bearing ϕ_cue_ and the vision based HD estimate α_vis_ that is defined by the anchoring angle γ_anchor_:

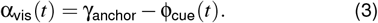

Note that we omit the notation for handling the periodicity of angles for simplicity. In reality, whenever we perform addition/subtraction of two angular measurements, we wrap this computation in an exponential and take the argument of the resulting exponential as a result.

The egocentric viewing angle ϕ_cue_ is defined as the difference between the true head direction α_true_ and the allocentric cue direction θ_cue_:

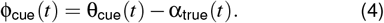

Substituting Eq. 4 in Eq. 3, we obtain a linear relationship (up to the angular periodicity) between the decoding error β_vis_ and the position dependent allocentric cue angle θ_cue_ (Fig.1c,d.):

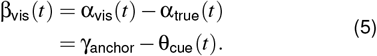

Intuitively, we can view the allocentric cue direction θ_cue_ as the angle of the vector from the subject to the cue. In this view, the linear relationship in Eq. 5 tells us that we expect a systematic, linear relationship between the HD bias and the position of the subject in the simple cue anchoring model. If the subject were only guided by visual input, we would expect that linear relationship to have a slope of − 1 and an offset given by the fixed anchoring angle.

To test whether this HD bias relation is also detectable in experimental data, we used the measured HD bias β_dec_ of each mouse as the angular differences between the tracked and internal HD. We also computed the allocentric cue angle θ_cue_(*t*) from the animals’ tracked position. We then performed a bootstrapped linear regression:

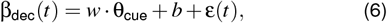

where the slope *w* and intercept *b* are fitted parameters minimizing the squared error of the linear fit to the observed data points. ε is the residual.

Eq. 6 corresponds to Eq 5, where the slope is *w* =− 1 and the intercept *b* =− *w ·* γ_anchor_. Note that the linear regression fits the y-intercept, while the anchoring angle is the x-intercept.

We use a linear regression between the decoding error β and the allocentric cue angle θ_cue_ to measure the amount of parallax and determine the anchoring angle. To account for autocorrelations in the time series data, we use stationary block bootstrapping (Politis & Romano, 1994) with an average block length of 2.5 s and n=1000 iterations to estimate the slopes, intercepts, and standard errors of *w* and *b*, as well as the p-values for the respective fits.

We apply the same procedure to obtain slopes and offsets for each entire session (after applying the rejection criteria described above), and for subsets of the session where the animal is looking directly towards or away from the cue to obtain the detected parallax in the “all data”, “in-view”, and “out-of-view” conditions.

### Computing the anchoring angle

The anchoring angle γ_anchor_ is calculated from the results of the bootstrapped linear regression as

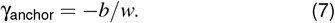

Uncertainties were propagated using standard error propagation:

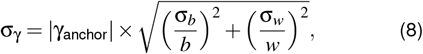

where σ_*b*_ and σ_*w*_ are the bootstrapped standard errors of the intercept and slope, respectively.

### Determining field of vision and weighting from cue switches

The integrate-and-vision model (Eq. 1) is dependent on a weighting factor σ and the field of vision. Both parameters can be estimated by analyzing the cue rotation periods of the single cue experiment recordings. At the beginning of each trial of these sessions, we observed a reorientation event, where the decoded HD took some time to adjust to the new cue’s position. We estimated this period by fitting a delayed exponential decay to the decoding error in these periods (Fig. 4e, purple). The time constant of the exponential decay τ tells us how fast the visual input changes the current HD estimate and is directly related to the weighting of the integrated and the visual input by 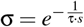, where *s* is the sampling rate. As we found the mean time constant is τ = 3.5 *s* (Fig. 4e), we obtained a weighting factor σ = 0.997.

Furthermore, we hypothesize that the delay before the onset of the exponential decay might be due to the effective field of vision of the animal, that is, the viewing angle range in which the animals are using cues for the visual anchoring. Based on the fitted anchoring angle, we observed that an effective field of vision of 180° best fits the fitted time delays (Fig. 4e-g).

### Software

Code was written in Python 3.12 using numpy, matplotlib, and scipy for analysis and visualization. Fits were performed using the scipy.stats.linregress function, bootstrapping using the arch package, and MSE minimization for σ using the optuna package.

## Data availability

For this work, the data from Duszkiewicz et al. (2024) and additional recordings in the same setting were used. The published data can be found here: 10.6084/m9.figshare.24921252. The additional data will be made available at the time of publication of this paper.

## Code availability

The code for the analysis and model will be made available at the time of publication of this paper.

## Acknowledgements

We thank Johannes Leugering, Melika Payvand, Paxon Frady and Viet Anh Khoa Tran and the memebers of the Wood/Dudchenko laboratory for their feedback. Generative AI was used, with care, to optimize and document code and to support writing. This research is funded by VolkswagenS-tiftung [CLAM 9C854], NIH [1R01EB026955] and the Simons Foundation [903332].

## Author contributions

A.R. and S.K. conceived and designed the study and the computational models. A.J.D. collected and processed the experimental data. S.K. performed the data analyses and implemented the models. A.R., E.N., and F.T.S. supervised the project. A.R., S.K., A.J.D., E.N., and F.T.S. wrote the manuscript.

## Supplementary Material

### Cue bearing vs. Cue azimuth

As this paper makes use of angle definitions in both allocentric and egocentric coordinate systems, it is important to clarify the nomenclature used here. In this work, we define east direction (x-axis / horizontal line) as the allocentric origin that represents zero degrees. All angles are measured counterclockwise. When using signed angular values, a positive angle is directed counterclockwise, and a negative angle is directed clockwise. Azimuth angles (also referred to as “directions”) are always defined as allocentric angles to the defined origin. So when we refer to the “cue azimuth”, this angle describes the allocentric angle of the vector between the animal and the cue (Suppl. Fig. 6, red). Note that this angle is only dependent on the position of the animal and the cue, and not on the animal’s head direction.

**Figure 6.**
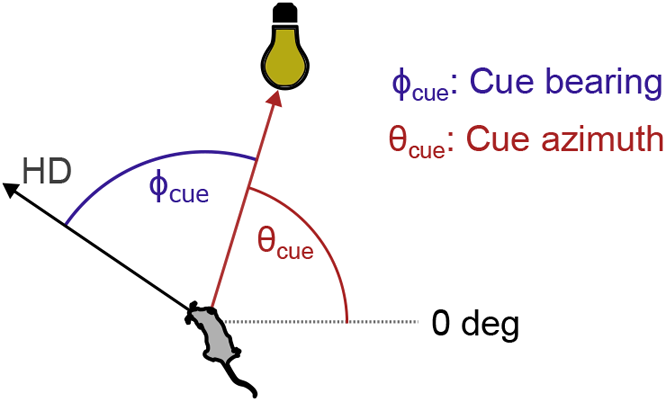
Diagram of the cue bearing and cue azimuth angles.

We use the word “bearing” to describe egocentric angles, that is, angles, that are measured relative to the current head direction. The cue bearing describes the angle between the current head direction and the cue azimuth (Suppl. Fig. 6, blue). Note that this angle is both dependent on the position of the animal and the cue, as well as the animal’s current head direction. The bearing angle can be understood as the angle at which the animal perceives a given object (e.g., there is a cue towards the right).

### Analysis of tuning curves from only the center of the environment

The analysis presented in the main part of this paper relies on the computed tuning curves from the neural activity. To ensure that there is no bias in the tuning curve that would explain the detected HD bias, we repeated the analysis, but computed the tuning curves only from time periods where the animal was in the center of the environment, where we expect the parallax to be minimal and therefore not influence the tuning curves. We found that the parallax effect remains detectable in this analysis and noted no qualitative differences from the analysis presented in the main text.

As can be seen in Fig. 7, we obtain the same detectable, but reduced parallax in the single cue setting and a strongly reduced parallax effect in the multi-cue setting. The cue visibility pattern also remains, i.e., we find a stronger parallax effect in the subset of time in each session where the animal is facing towards the cue and a lower amount of parallax in time periods where the animal is facing away from the cue.

**Figure 7.**
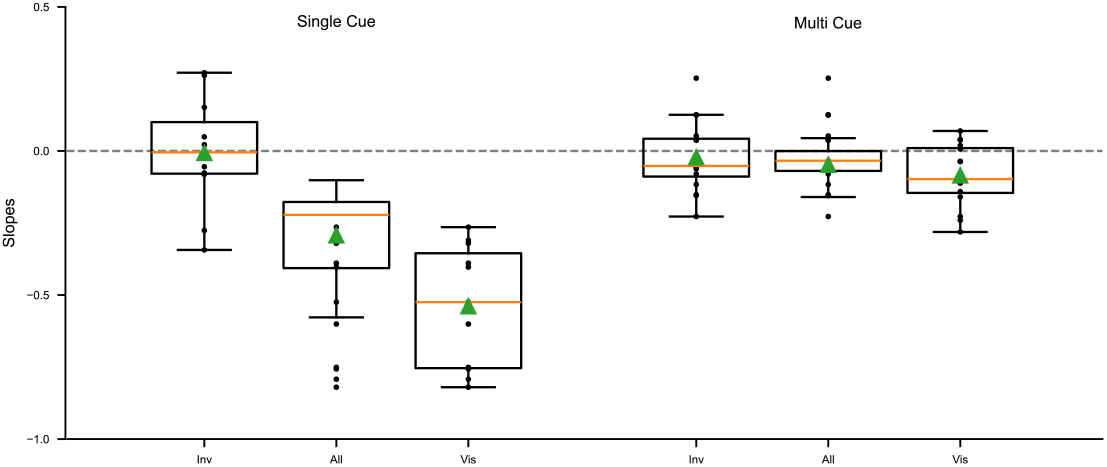
Results of parallax analysis using tuning curves from the center of the environment. The distribution of measured slopes from the linear regression is presented in the two experimental setups, “Single cue” and “multi-cue”. The three conditions “in-view”, “all data”, and “out-of-view” (from left to right) are defined as in the main text (see Figs. 1h inset and 3d,h)

**Figure 8.**
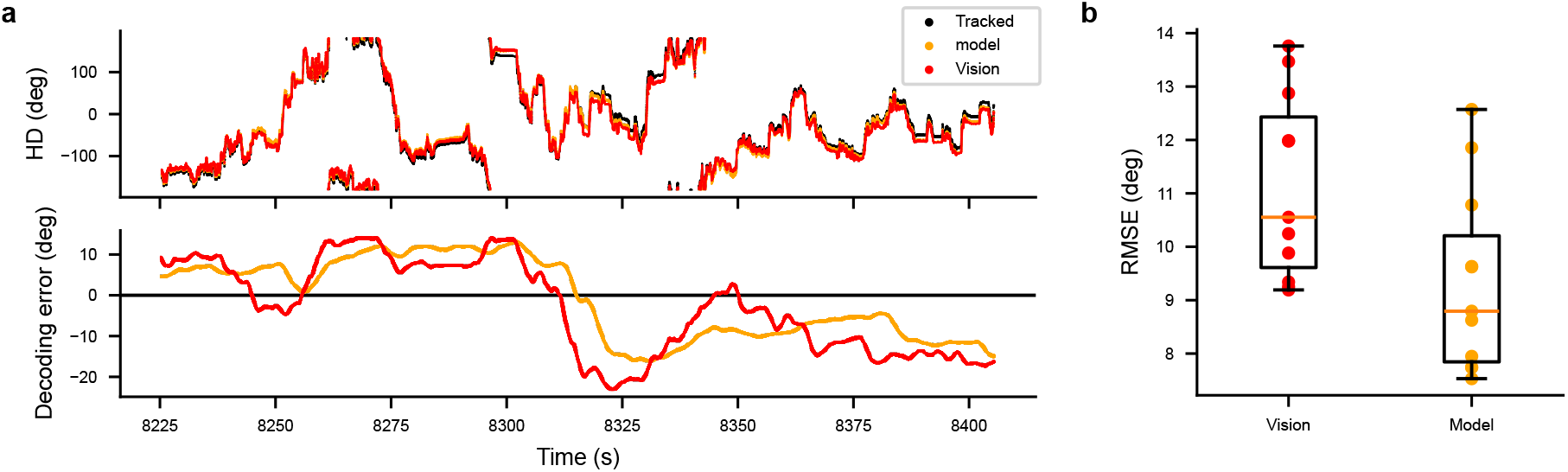
AHV integration shows reduced HD tracking error compared to a purely vision-based estimation. **a:** Head direction over time in an example simulation. Top panel: Tracked HD (black) indicates the ground truth in the simulation. Vision-based HD (red) resembles the estimate of a purely vision-based estimation of the head direction from a single cue (full parallax). The yellow trace indicates the results from the “integrate-and-vision” model, where both visual estimates and AHV integrated estimates are combined at each time step. Bottom panel: Residual plot of the HD tracking error over time for the full model (yellow) and the purely vision-based estimation (red). **b:** The root mean squared error (RMSE), i.e., the mean distance between the black and the yellow/red traces for all simulations. across all simulated sessions for the purely vision based (left) estimate and the model estimate (right)

**Figure 9.**
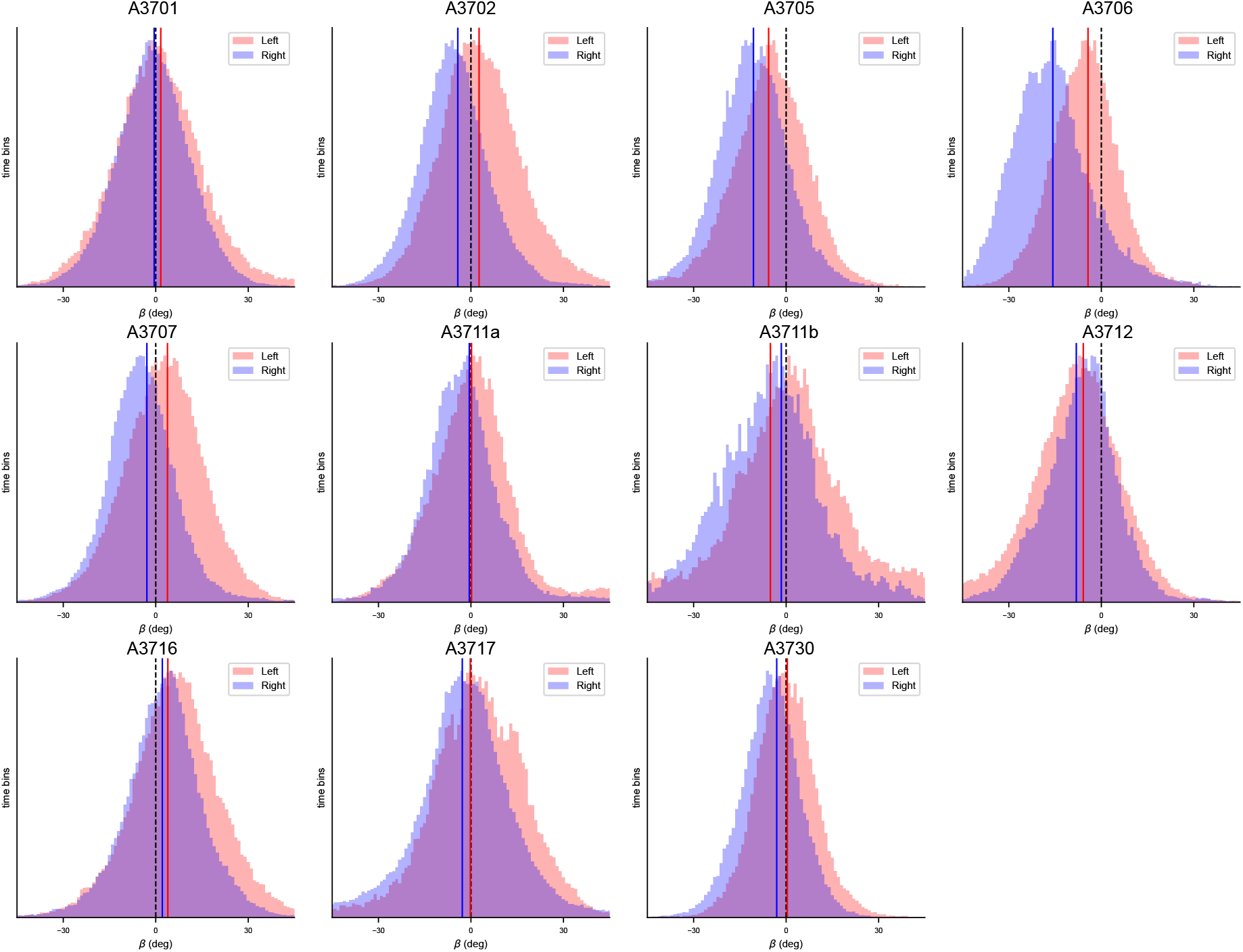
Decoding error histogram split by left and right region in environment for single cue datasets.

**Figure 10.**
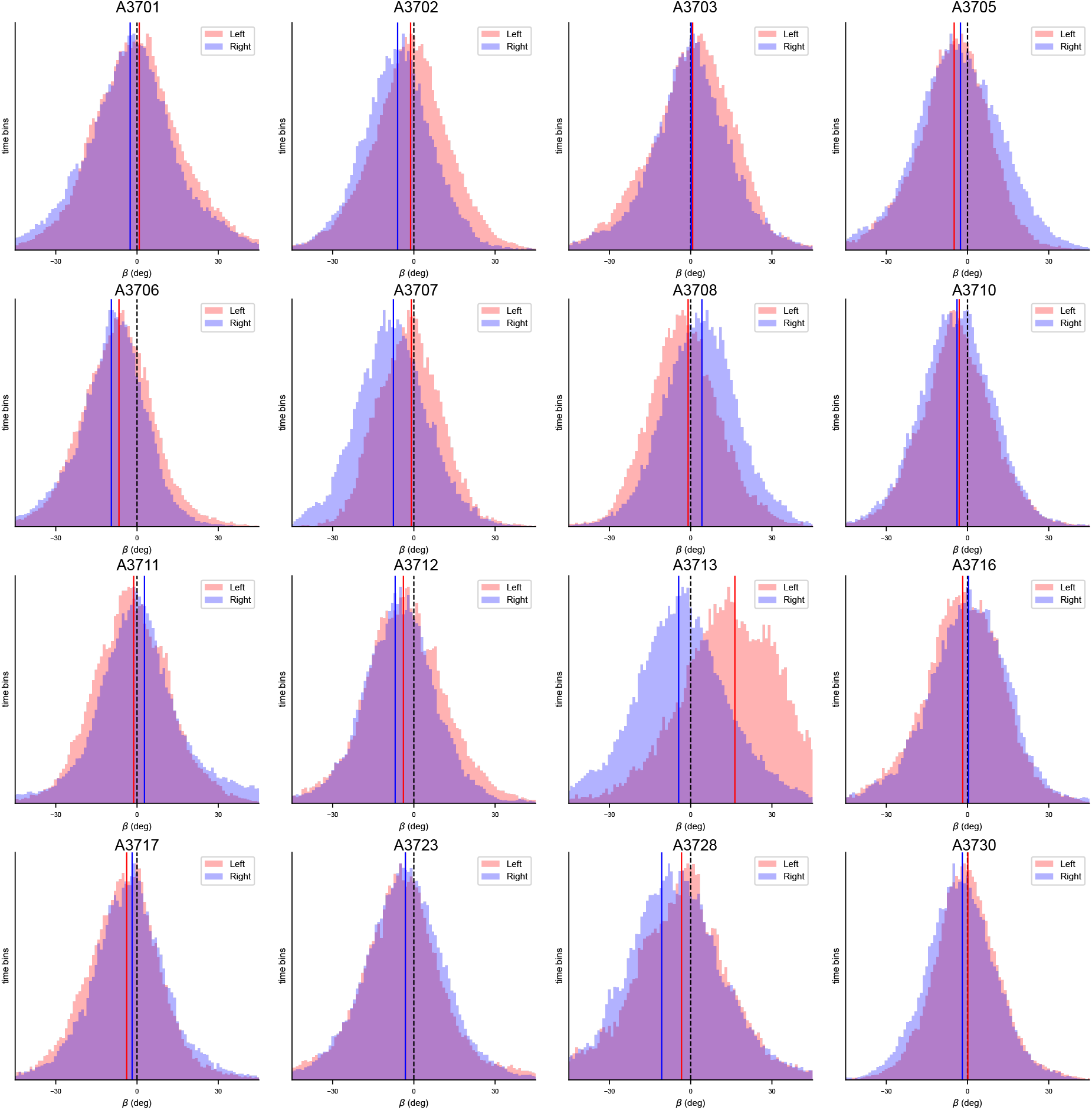
Decoding error histogram split by left and right region in environment for multi-cue dataset.

**Figure 11.**
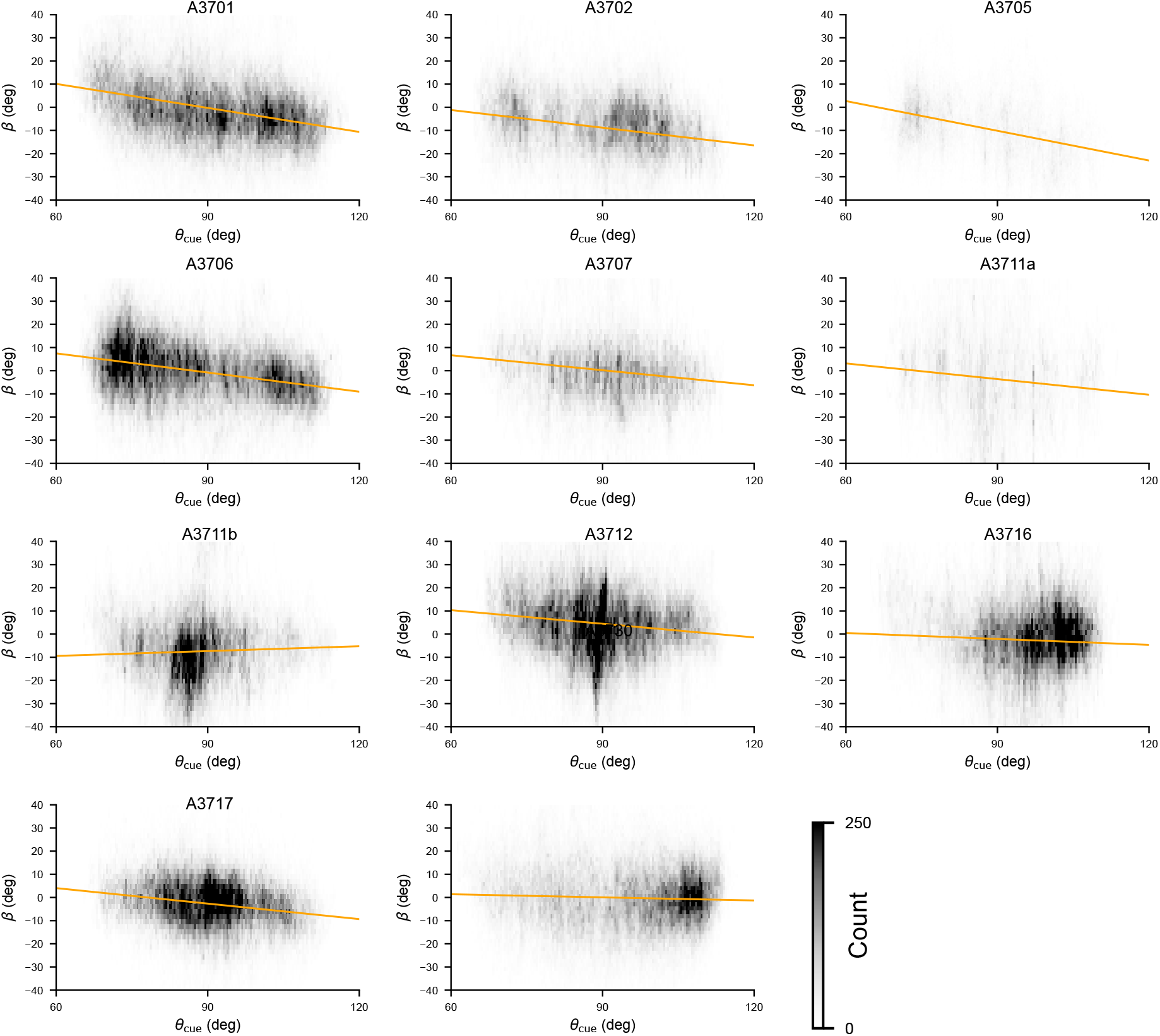
Linear regression for the single cue dataset.

**Figure 12.**
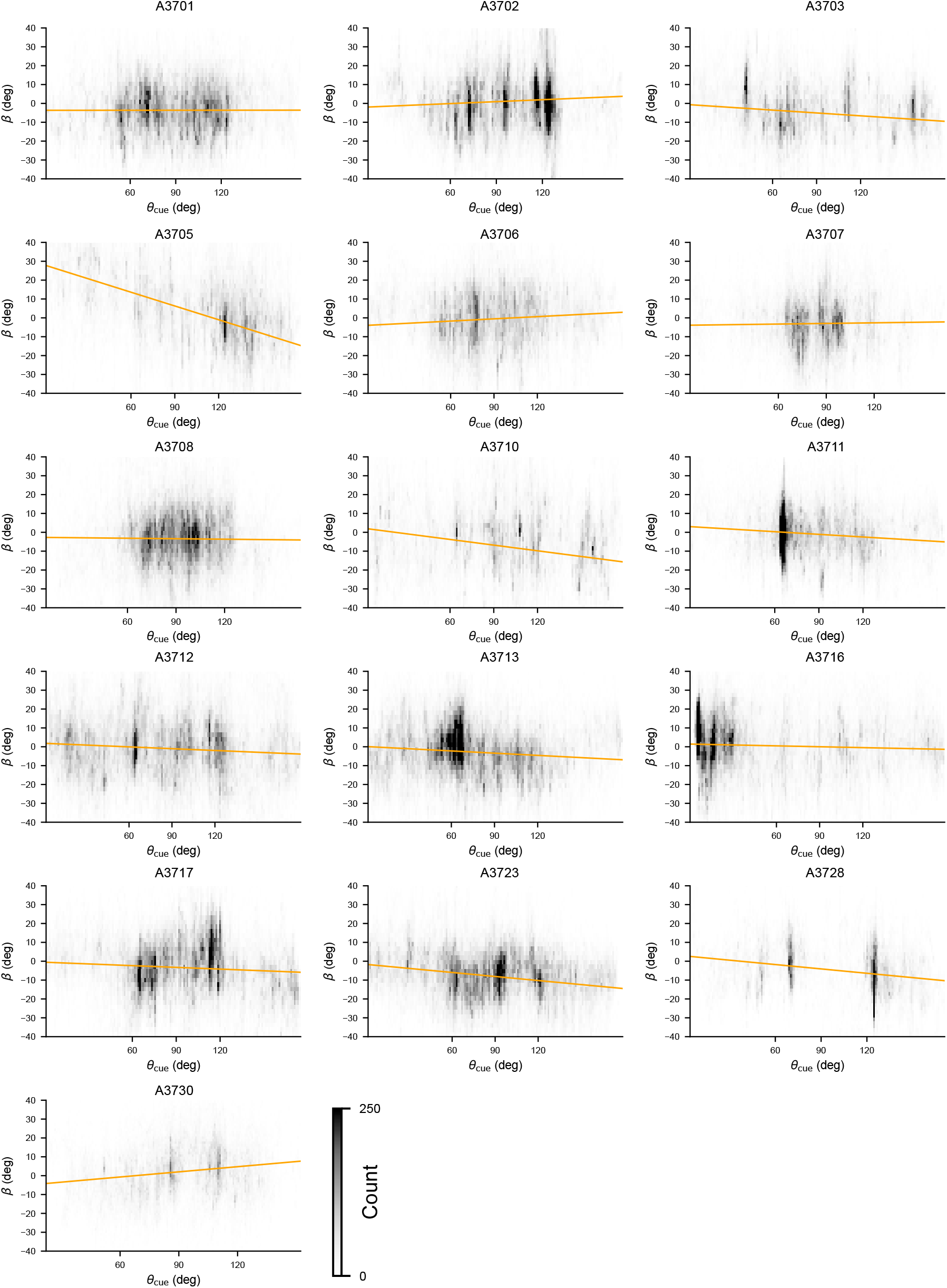
Linear regression for the multi-cue dataset.

**Figure 13.**
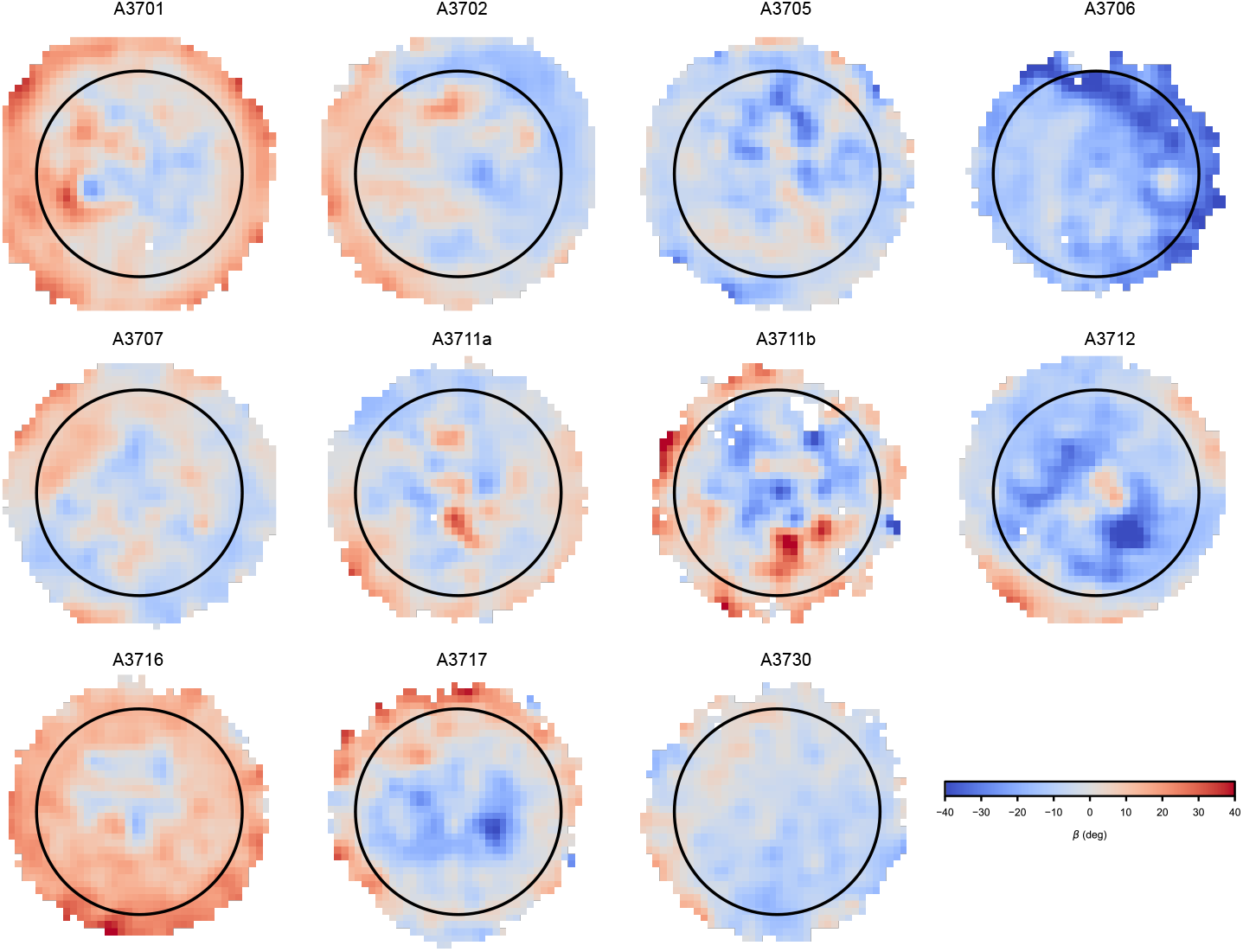
Heat maps of measured parallax across space. Same as Fig. 2, here for all sessions in the single cue dataset.

**Figure 14.**
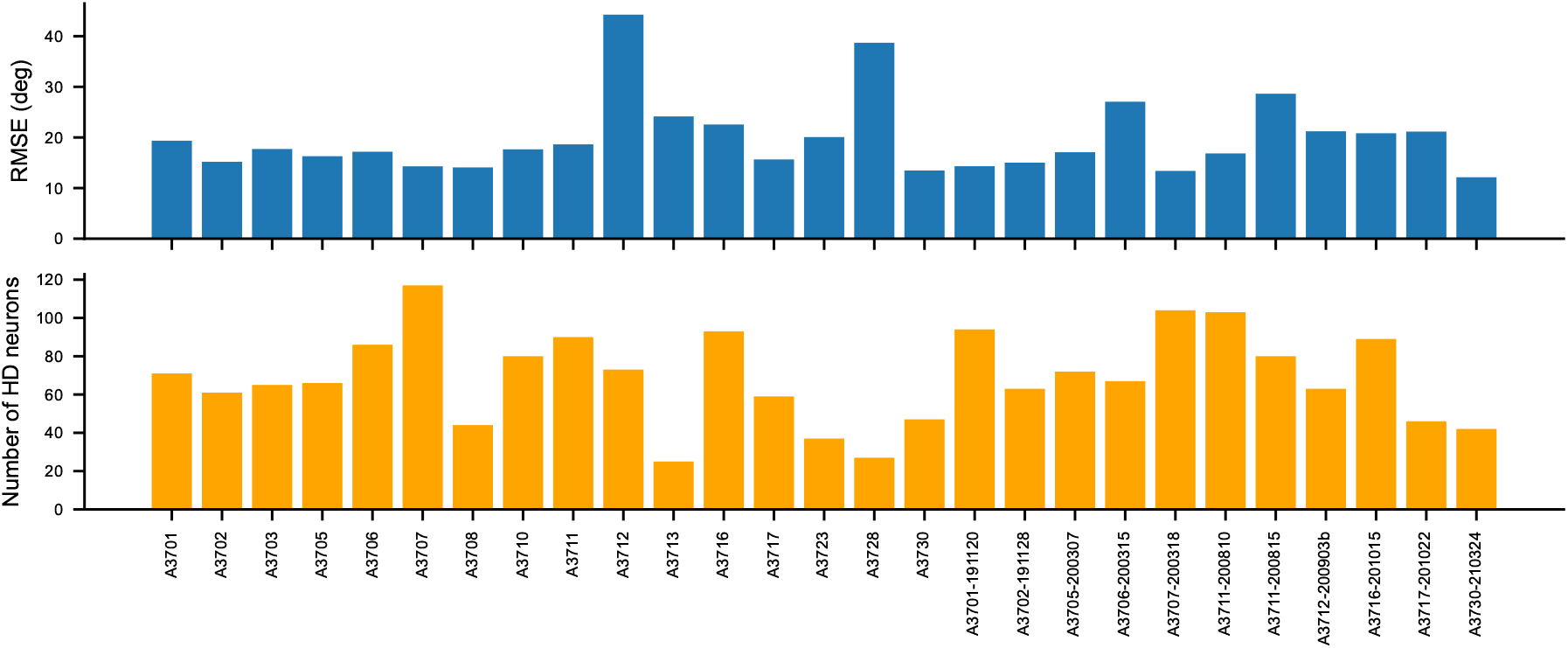
Mean decoding error and number of neurons per session. **(a)**: Bars show the root mean squared error between the tracked and decoded head direction. Switch periods and other rejected periods due to tracking errors are excluded from this data. **(b)**: Number of neurons recorded in each session.

### AHV integration improves HD tracking in simulation

To illustrate the improved HD tracking accuracy when including the AHV integration, we compare the “integrate-and-vision” model (Sec. Simple integration passively attenuates parallax error) to a purely vision-based estimation of the head direction in the single-cue setting. The inclusion of AHV integration in the modeled HD estimate attenuates the full parallax effect, producing a more accurate estimate of the head direction in our simulations (Suppl. Fig. 8).

The use of the AHV integration results in a lower root mean squared error (RMSE) across all simulated sessions. The example trace shown in Suppl. Fig. 8 highlights how the inclusion of AHV integration reduces the influence of the parallax effect from the pure vision model, leading to smaller errors in tracking the head direction.

## Significance testing results

**Table 1:** Head Direction Bias Analysis Results.

| Session | Left Side |  |  | Right Side |  |  | Statistics |  |  |
| --- | --- | --- | --- | --- | --- | --- | --- | --- | --- |
| | $n_{\text{eff}}$ | Mean ($^{\circ}$) | SD ($^{\circ}$) | $n_{\text{eff}}$ | Mean ($^{\circ}$) | SD ($^{\circ}$) | $t(\text{DoF})$ | $p$ | $\vert d\vert $ |
| A3701-191120 | 611.2 | 1.64 | 16.27 | 1599.7 | -0.50 | 12.70 | 2.93(908.4) | 0.003** | 0.15 |
| A3702-191128 | 678.0 | 2.64 | 14.86 | 905.8 | -4.26 | 13.05 | 9.63(1347.9) | < .001*** | 0.49 |
| A3705-200307 | 1362.1 | -5.68 | 14.26 | 3311.0 | -10.60 | 14.12 | 10.75(2511.4) | < .001*** | 0.35 |
| A3706-200315 | 389.8 | -4.32 | 10.33 | 223.9 | -15.73 | 12.83 | 11.36(388.9) | < .001*** | 1.00 |
| A3707-200318 | 1126.0 | 3.81 | 13.62 | 1917.5 | -2.88 | 12.81 | 13.35(2241.4) | < .001*** | 0.50 |
| A3711-200810 | 343.1 | 0.20 | 12.83 | 1425.9 | -0.54 | 16.71 | 0.90(651.9) | 0.371 | 0.05 |
| A3711-200815 | 192.0 | -5.10 | 25.35 | 2005.3 | -1.56 | 29.30 | -1.82(242.7) | 0.070 | 0.13 |
| A3712-200903b | 1303.8 | -5.85 | 14.44 | 9717.1 | -8.12 | 20.41 | 5.04(2074.8) | < .001*** | 0.12 |
| A3716-201015 | 6577.6 | 3.93 | 17.89 | 5898.4 | 2.14 | 16.06 | 5.88(12474.0) | < .001*** | 0.11 |
| A3717-201022 | 885.4 | -0.21 | 19.28 | 3898.1 | -2.81 | 20.76 | 3.58(1389.9) | < .001*** | 0.13 |
| A3730-210324 | 1218.8 | 0.37 | 11.31 | 944.5 | -3.05 | 10.25 | 7.37(2109.0) | < .001*** | 0.32 |

**Table 2:** One-Sided, one-sample t-tests testing whether slope distributions significantly differ from zero. Significance levels:*** *p* < 0.001, ** *p* < 0.01, * *p* < 0.05, ns = not significant.

| Dataset | Condition | Mean slope | SD | t(DoF) | p-value | Significance |
| --- | --- | --- | --- | --- | --- | --- |
| Single Cue |  |  |  |  |  |  |
| | In-view | -0.508 | 0.216 | $t(10) = -7.79$ | $< 0.001$ | *** |
| | All data | -0.202 | 0.139 | $t(10) = -4.83$ | $< 0.001$ | *** |
| | Out-of-view | 0.002 | 0.195 | $t(10) = 0.04$ | 0.515 | ns |
| Multi-Cue |  |  |  |  |  |  |
| | In-view | -0.069 | 0.116 | $t(15) = -2.38$ | 0.015 | * |
| | All data | -0.034 | 0.074 | $t(15) = -1.83$ | 0.044 | * |
| | Out-of-view | -0.010 | 0.118 | $t(15) = -0.35$ | 0.365 | ns |

All tests used one-sided, one-sample t-tests against zero after Shapiro-Wilk normality testing.

**Table 3:**
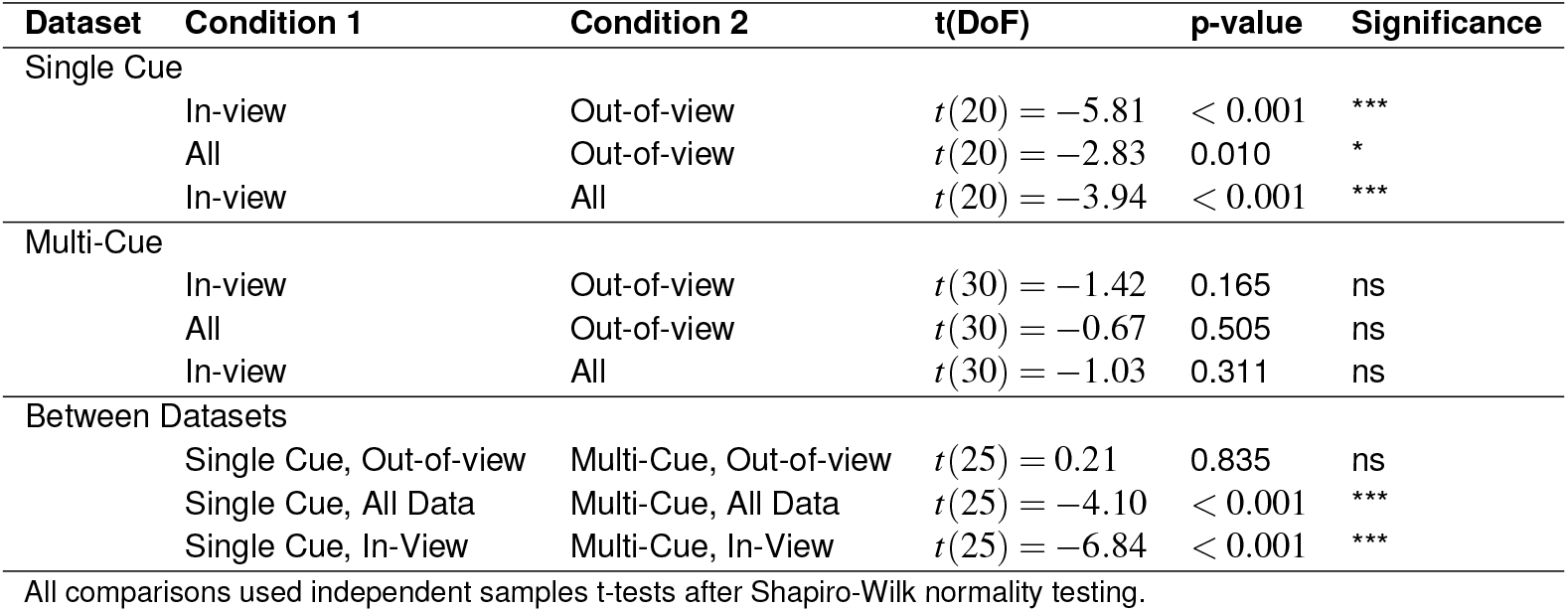
Statistical comparison of slope distributions across viewing conditions and datasets. Slopes represent the relationship between landmark angle and decoding error. Significance levels: *** *p* < 0.001, ** *p* < 0.01, * *p* < 0.05, ns = not significant.

**Table 4:**
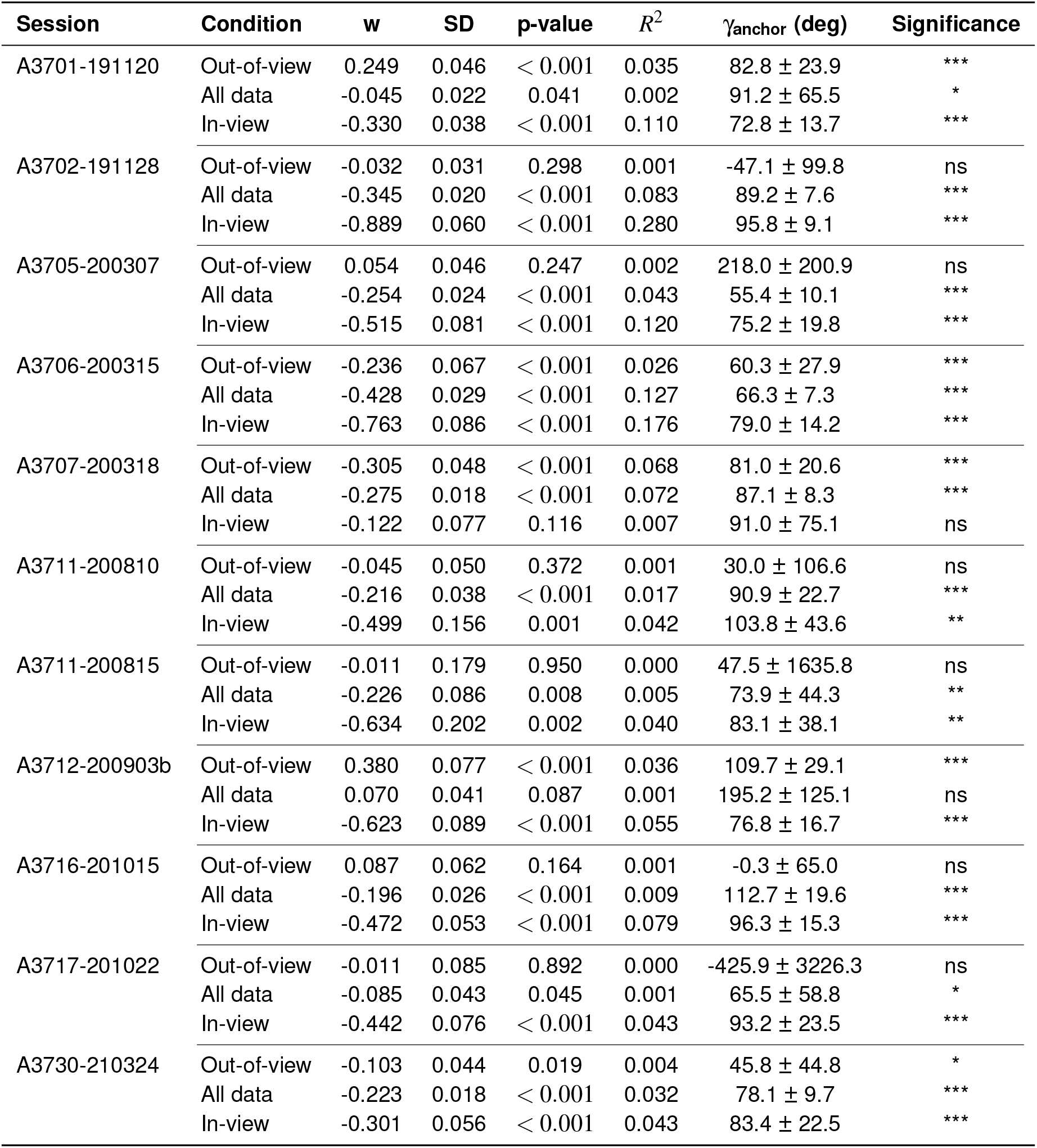
Parallax Analysis Results - Single Cue. Anchoring angles with errors > 180° are not reported.

**Table 5:** Parallax Analysis Results - Multi Cue. Anchoring angles with errors > 180° are not reported.

| Session | Condition | w | SD | p-value | $R^2$ | $\gamma_{\text{anchor}}$ (deg) | Significance |
| --- | --- | --- | --- | --- | --- | --- | --- |
| A3701 | Out-of-view | -0.090 | 0.017 | $< 0.001$ | 0.045 | $49.5 \pm 20.4$ | *** |
| | All data | -0.032 | 0.011 | 0.003 | 0.005 | $58.2 \pm 38.2$ | ** |
| | In-view | 0.127 | 0.027 | $< 0.001$ | 0.061 | $83.5 \pm 29.1$ | *** |
| A3702 | Out-of-view | -0.013 | 0.012 | 0.313 | 0.001 | $-462.4 \pm 470.7$ | ns |
| | All data | -0.039 | 0.010 | $< 0.001$ | 0.010 | $3.9 \pm 23.3$ | *** |
| | In-view | -0.044 | 0.023 | 0.052 | 0.012 | $95.8 \pm 65.4$ | ns |
| A3703 | Out-of-view | -0.034 | 0.011 | 0.002 | 0.013 | $95.6 \pm 41.0$ | ** |
| | All data | -0.016 | 0.008 | 0.054 | 0.002 | $90.8 \pm 63.6$ | ns |
|  | In-view | 0.002 | 0.028 | 0.955 | 0.000 | - | ns |
| A3705 | Out-of-view | 0.062 | 0.024 | 0.010 | 0.016 | $135.9 \pm 64.6$ | ** |
| | All data | -0.030 | 0.012 | 0.011 | 0.005 | $-17.1 \pm 39.8$ | * |
| | In-view | -0.194 | 0.019 | $< 0.001$ | 0.210 | $75.9 \pm 11.6$ | *** |
| A3706 | Out-of-view | -0.068 | 0.022 | 0.002 | 0.027 | $-38.5 \pm 26.7$ | ** |
| | All data | -0.071 | 0.010 | $< 0.001$ | 0.034 | $-23.4 \pm 13.2$ | *** |
| | In-view | -0.138 | 0.016 | $< 0.001$ | 0.111 | $42.4 \pm 12.6$ | *** |
| A3707 | Out-of-view | -0.061 | 0.027 | 0.024 | 0.015 | $-5.6 \pm 57.4$ | * |
| | All data | -0.077 | 0.014 | $< 0.001$ | 0.051 | $37.2 \pm 20.5$ | *** |
| | In-view | -0.224 | 0.034 | $< 0.001$ | 0.288 | $61.5 \pm 15.7$ | *** |
| A3708 | Out-of-view | 0.132 | 0.027 | $< 0.001$ | 0.094 | $71.2 \pm 23.7$ | *** |
| | All data | 0.091 | 0.019 | $< 0.001$ | 0.027 | $67.8 \pm 23.8$ | *** |
| | In-view | -0.213 | 0.065 | 0.001 | 0.044 | $104.9 \pm 43.7$ | ** |
| A3710 | Out-of-view | 0.007 | 0.023 | 0.749 | 0.000 | $689.9 \pm 2179.9$ | ns |
|  | All data | 0.000 | 0.014 | 0.974 | 0.000 | - | ns |
| | In-view | -0.059 | 0.019 | 0.002 | 0.012 | $68.3 \pm 37.4$ | ** |
| A3711 | Out-of-view | 0.030 | 0.025 | 0.233 | 0.003 | $115.7 \pm 127.9$ | ns |
| | All data | 0.034 | 0.020 | 0.086 | 0.003 | $62.8 \pm 62.6$ | ns |
| | In-view | 0.075 | 0.040 | 0.060 | 0.012 | $48.4 \pm 55.4$ | ns |
| A3712 | Out-of-view | 0.073 | 0.039 | 0.064 | 0.007 | $190.0 \pm 113.7$ | ns |
| | All data | -0.051 | 0.019 | 0.008 | 0.002 | $-10.0 \pm 39.0$ | ** |
| | In-view | 0.004 | 0.052 | 0.937 | 0.000 | $52.9 \pm 1249.9$ | ns |
| A3713 | Out-of-view | -0.264 | 0.024 | $< 0.001$ | 0.242 | $112.9 \pm 14.5$ | *** |
| | All data | -0.243 | 0.013 | $< 0.001$ | 0.187 | $115.9 \pm 8.2$ | *** |
| | In-view | -0.248 | 0.025 | $< 0.001$ | 0.211 | $125.2 \pm 18.1$ | *** |
| A3716 | Out-of-view | 0.192 | 0.059 | 0.001 | 0.043 | $106.4 \pm 45.9$ | ** |
| | All data | 0.039 | 0.018 | 0.030 | 0.004 | $103.5 \pm 67.1$ | * |
| | In-view | -0.025 | 0.028 | 0.372 | 0.002 | $113.3 \pm 162.5$ | ns |
| A3717 | Out-of-view | -0.012 | 0.026 | 0.649 | 0.001 | $-406.9 \pm 913.6$ | ns |
| | All data | 0.011 | 0.018 | 0.532 | 0.000 | $359.7 \pm 592.8$ | ns |
| | In-view | 0.088 | 0.045 | 0.049 | 0.011 | $130.0 \pm 79.3$ | * |
| A3723 | Out-of-view | 0.133 | 0.039 | $< 0.001$ | 0.027 | $127.7 \pm 44.8$ | *** |
| | All data | -0.009 | 0.023 | 0.705 | 0.000 | $-305.9 \pm 848.8$ | ns |
| | In-view | -0.151 | 0.031 | $< 0.001$ | 0.029 | $80.2 \pm 26.5$ | *** |
| A3728 | Out-of-view | -0.190 | 0.042 | $< 0.001$ | 0.057 | $25.4 \pm 19.6$ | *** |
| | All data | -0.100 | 0.024 | $< 0.001$ | 0.015 | $22.3 \pm 25.5$ | *** |
| | In-view | -0.075 | 0.081 | 0.354 | 0.006 | $116.3 \pm 166.7$ | ns |
| A3730 | Out-of-view | -0.063 | 0.026 | 0.014 | 0.018 | $88.6 \pm 60.4$ | * |
| | All data | -0.047 | 0.011 | $< 0.001$ | 0.012 | $67.0 \pm 26.8$ | *** |
| | In-view | -0.032 | 0.023 | 0.163 | 0.004 | $73.4 \pm 75.4$ | ns |

## Supplementary plots for all sessions

In this section, we provide additional plots for all sessions in cases where we only showed example sessions in the main text.

